# Genomic Population Structure of the Sunda Pangolin (*Manis javanica*) in Thailand: Implications for Conservation and Wildlife Forensics

**DOI:** 10.1101/2025.10.06.680665

**Authors:** Nattapong Banterng, Kyle Ewart, Rob Ogden

## Abstract

Pangolins are the most heavily trafficked mammal globally, yet the population structure and genetic diversity remain poorly understood for many of the eight species. In Thailand, the Sunda pangolin (*Manis javanica*) is subject to illegal hunting and has a fragmented distribution among protected areas; however, relatively little is known about its genetic variation, due in part to the elusive nature of the species, which presents significant challenges for sampling and population genetic research. This study analyses whole genome sequences from 30 georeferenced samples across Thailand to examine population structure and genetic diversity, in order to inform conservation management and underpin the development of wildlife forensic tools. We identified four distinct genetic populations within Thailand, corresponding to the Western, Khao Yai, Mid-south, and Southernmost regions, with moderate genetic differentiation (*F_ST_* = 0.06-0.14) between populations. The Western population showed elevated inbreeding levels (*F_ROH_* = 18.50%) compared to other populations (*F_ROH_* < 12.5%), while the Southernmost population maintained the highest genetic diversity. Geographic barriers, including the Chao Phraya River and Nakhon Si Thammarat Mountain range, coincided with strong population structure. Our findings establish a foundational genomic baseline for Sunda pangolin conservation in Thailand, identifying distinct genetic management units. The population-specific genetic signatures provide a framework for developing wildlife forensic tools to trace the geographic origin of confiscated pangolins within Thailand. These insights are crucial for implementing targeted conservation strategies and combating illegal wildlife trade of this Critically Endangered species.

## Introduction

Within wildlife conservation research, the use of whole-genome sequence data for population genomic studies is providing greater insights into genetic diversity and population structure than ever before (Hohenlohe, Funk, and Rajora 2021; Barbosa et al. 2021). Such studies rely on samples of known geographic provenance to serve as essential references for understanding biogeographical processes (Forester et al. 2021) and tracing the origins of individuals, particularly in the context of illegal wildlife trade (Ogden and Linacre 2015). In the context of conservation genomics, such data can inform management strategies by delineating conservation units, identifying ecological niches and prioritising areas for protection that ensure the long-term persistence of endangered species (Funk et al. 2012; Coates, Byrne, and Moritz 2018). Moreover, genomic data with geographical references have been instrumental in identifying source populations, trafficking routes and trade hubs for various trafficked species (Presti et al. 2015; Wasser et al. 2022), including species such as the white-bellied pangolin *(Phataginus tricuspis)* (Tinsman et al. 2023).

Pangolins are the most heavily trafficked mammals in the world (Challender et al. 2020), with trade propelled by demand for their scales for traditional medicines, and their meat for consumption as a gastronomic delicacy (Wong 2019). Within Asia, the Critically Endangered Sunda pangolin, *Manis javanica*, ranges from pristine ecosystems to urban landscapes across Southeast Asia and southern China, yet population data for this species is scarce (Challender et al. 2019; Chong et al. 2020). Evidence indicates a downward trend in population numbers throughout significant portions of its distribution due to overexploitation and trade at the local and international level, resulting in the species being listed on CITES Appendix I (Chong et al. 2020). Previous studies have shown the utility of genetic analysis of scale seizures to identify Sunda pangolins (Ewart et al. 2021; Priyambada et al. 2021) and to infer their geographic origin(s) (Sitam et al. 2023; Nash et al. 2018). However, a lack of extensive geo-referenced samples from the species limits our knowledge of population structure and diversity necessary for precise traceability testing (Zhang et al. 2020; Gu, Hu, and Yu 2024). This scarcity of reference data is largely due to the species’ ecology - Sunda pangolins live in woodland burrows and are largely nocturnal (Chong et al. 2020), making studies of their ecology or population genetic structure notoriously difficult and severely limiting sample sizes, even prior to anthropogenic population declines (Heighton and Gaubert 2021; Gu, Hu, and Yu 2024; Shirley et al. 2023). The use of whole genome sequencing and the associated extensive genetic marker panels can mitigate this issue to some extent, by enabling characterisation of genetic variation at a locality using fewer samples than were required for traditional population genetic approaches (Allendorf, Hohenlohe, and Luikart 2010; Flanagan et al. 2018; Theissinger et al. 2023).

Previous population genetic studies of the Sunda pangolin revealed considerable genetic structure, with the northern Chinese and Myanmar populations differentiated from other Southeast Asia populations (Hu et al. 2020), the clustering of populations from Borneo, Java and Singapore/Sumatra (Nash et al. 2018), and population differentiation within Borneo (Sitam et al. 2023). However, there remains a paucity of reference data for this species (Zhang et al. 2015; Sitam et al. 2023), hence the picture of population structure throughout the species’ range in incomplete. Within Thailand, there is evidence for the existence of multiple evolutionary lineages separating the eastern, western and southern parts of the Sunda pangolin distribution, based on whole mitogenome sequence data (Banterng et al. 2025), but a lack of nuclear genetic marker data hinders our understanding of contemporary population structure and gene flow in this region and prevents robust genetic geographic origin assignment.

In this context, the Thai Department of National Parks, Wildlife and Plant Conservation identified the need for more precise analysis of Sunda pangolin populations. In response, this study aimed to apply whole genome sequencing of geo-referenced samples throughout Thailand to investigate the genetic diversity and population structure of Sunda pangolins. These data will inform conservation management of the species and provide a genetic baseline for tracing the origins of confiscated Sunda pangolin individuals at a fine geographic scale.

## Materials and Methods

### Ethics statement, permissions and permitting

This study was conducted in compliance with the Animal Research: Reporting of In Vivo Experiments (ARRIVE) guidelines for animal research, with ethical approval to collect the samples received from the University of Edinburgh Animal Welfare and Ethical Review Board (AWERB) (No. OS2-22). All samples were taken from wild pangolins, either dead or alive, from inside and outside-protected areas under permission from the Department of National Parks Wildlife and Plant Conservation, Thailand (Permit No.0909.204/9311). Sampling was carried out in accordance with the Wildlife Protection and Preservation Act of 2562 B.E. and relevant regulations. No pangolins were sacrificed for this study; blood samples were collected under permit, without anaesthesia. Samples were imported to the University of Edinburgh, UK, from Thailand under CITES export permit No. 22TH0902.2/521 and 23TH0902.2/2 (Thailand) and the CITES import permit No. 616968/01-03 and 624109/01 (UK).

### Collection and DNA extraction

Sunda pangolin blood (*n = 25*) and tissue (*n = 6*) samples were collected opportunistically from rescued (*n = 30*) and confiscated (*n = 1*) pangolins of known geographic origin across Thailand (*Sup. Table S1*). The geographical origin of the single confiscated specimen was based on enforcement records; the animal was seized from local hunters who were apprehended within specific protected areas, and who confirmed the collection location of the Sunda pangolin individual during their arrest. Given the nature of these enforcement operations, it is extremely unlikely that this specimen derived from long-distance trafficking. Blood samples were collected in EDTA tubes, and tissue samples were collected in plastic tubes and stored at −20°C before analysis. DNA was extracted using the Invitrogen PureLink™ Genomic DNA Mini Kit (Thermo Fisher Scientific, Carlsbad, CA, USA; Cat. No. K182001) following the manufacturer’s protocol. Existing Sunda pangolin sequence data were downloaded from NCBI GenBank (*Sup. Table S1*).

### Genome assembly and variant calling

The 31 DNA samples were subject to whole genome sequencing using the Illumina NovaSeq platform provided by Azenta Life Sciences (Takeley, Essex,UK). Paired-end 150 bp reads were generated to target an average sequencing depth of coverage of 10X across the genome. All paired-end reads were trimmed to remove low quality bases and adapter sequences using TrimGalore 0.6.6, with trimming parameters set to remove bases with Phred quality scores <30 and to discard reads shorter than 35 bp after trimming. The trimmed reads were then mapped against a Sunda pangolin reference genome (GCA_024605085.1); as this accession is limited to a contig-level assembly, we contacted the author to provide the constructed pseudo-chromosome versions as mentioned in the associated publication (Yan et al. 2022). Paired-end read data from NCBI generated from samples collected from Yunnan Province (China), Kachin State (Myanmar) and Malaysia were processed using the same method. As these five individuals lack precise location information, we set the location at the geographic mid-point of the province or state declared.

Trimmed sequence reads were mapped against the constructed pseudo-chromosome level reference genome with the BWA-MEM algorithm v0.7.17 using default settings (Li and Durbin 2009), and variants were called using DeepVariant v1.5.0 with --model_type: WGS, a model that is best suited for Illumina Whole Genome Sequencing data (Poplin et al. 2018), and then joint calling was performed using GLnexus v1.3.1 (Yun et al. 2021) with --config DeepVariantWGS, a configuration for DeepVariant output. The variant data files were filtering with PLINK v1.9 (Purcell et al. 2007), BCFtools v1.9 (Danecek et al. 2021) and VCFtools v0.1.13 (Danecek et al. 2011). We retained only biallelic SNPs, removed SNPs with >20% missing genotypes, excluded samples with >20% missing genotypes, and removed SNPs with a minor allele frequency (MAF) <5%. For analysis that required SNPS in linkage disequilibrium (LD), a sliding window approach in PLINK (window size = 50 SNPs, step size = 10 SNPs, r² threshold = 0.2) was carried out SNPs were pruned when r² > 0.2.

Given the small sample size and geographic differentiation of the Myanmar and China populations, ascertainment bias could be an issue. This was investigated through inspection of allele frequency spectra (Marth et al. 2004; Nielsen, Hubisz, and Clark 2004) for each putative genetic population using VCFtools.

As our samples were collected opportunistically from rescued or confiscated pangolins, to ensure that none of the samples included in the study were closely related to each other, pairwise relatedness between individuals was calculated by combining output from the PLINK --*genome* function in PLINK and estimated using the KING, R0 and R1 coefficients in NgsRelate v2 (Hanghøj et al. 2019).

### Population structure

We used a principal components analysis (PCA) within PLINK to investigate genetic clustering, followed by ADMIXTURE v1.3 (Alexander, Novembre, and Lange 2009) to estimate individual ancestry within a hypothetical number of the subpopulations (*K*), from *K* = 1-7. Following identification of putative genetic populations, we used VCFtools to calculate weighted (Weir-Cockerham) pairwise *F_ST_* to examine differentiation between the populations.

### Genetic diversity, inbreeding and ROH detection

VCFtools was used to calculate nucleotide diversity (π) in 20 Mb windows across the genome,and to estimate inbreeding coefficients (*F_IS_*) for each individual using the method of moments approach (Ritland 1996).

ANGSD v0.923 (Korneliussen, Albrechtsen, and Nielsen 2014) was used to estimate genome-wide heterozygosity for each sample following the ANGSD recommended pipeline. We implemented the *doSaf* command on the BAM alignment files to infer folded site allele frequency likelihoods. Quality control measures were applied to ensure reliable estimates: reads were filtered to remove those that failed platform/vendor quality checks (- remove_bads 1), did not map uniquely (-uniqueOnly 1), or lacked proper pair mapping (- only_proper_pairs 1). We implemented base alignment quality adjustment (-baq 1) and enforced minimum thresholds of Phred 20 for both mapping quality (-minMapQ 20) and base quality (-minQ 20). To account for coverage variation, we restricted analysis to sites with depths between 1 and 21 reads (-setMinDepth 1 -setMaxDepth 21).

The site allele frequency was estimated using the reference genome as the ancestral state, with genotype likelihoods calculated using -GL 2. The analysis was performed using a folded spectrum approach (-fold 1) to focus on heterozygosity estimation. Following the initial frequency estimation, we used the ANGSD subprogram *realSFS* to infer the site frequency spectrum for each sample. Statistical differences in heterozygosity among populations were tested using one-way ANOVA followed by pairwise t-tests with Bonferroni correction. We also tested the distribution of heterozygosity values for normality using Shapiro-Wilk tests for each population.

We identified runs of homozygosity (ROH) with a minimum length of 500 kb and at least 50 SNPs using the PLINK (Purcell et al. 2007) --homozyg function with the following parameters: --homozyg-window-snp 50, --homozyg-snp 50, --homozyg-kb 500, --homozyg-gap 1000, -- homozyg-density 50, --homozyg-window-missing 5, and --homozyg-window-het 3. Individual inbreeding, as the proportion of the genome in runs of homozygosity (*F_ROH_*), was calculated as the sum of the detected ROH lengths for each individual divided by the total autosomal assembly length (2443 Mb).

## Results

### Variant calling and relatedness analysis

One Thailand sample, X11, was removed from downstream analysis due to low sequence quality. From the remaining 30 whole genome datasets generated in this study, combined with the 5 genome datasets from NCBI, we retrieved 30,611,645 nucleotide variants. After filtering, this reduced to 15,940,461 biallelic SNPs and subsequently to 1,071,601 LD-pruned biallelic SNPs. The mean variant sequencing depth among individuals ranged from 9.8-19.2 (*Sup. Table S3*).

Analysis of allele frequency distributions among localities revealed that the China and Myanmar (CHMM) samples exhibited a strongly bi-modal distribution, with most alleles being at very low or very high frequencies, suggesting potential ascertainment bias affecting the results, (*Sup. Fig.S1*) (Marth et al. 2004; Lachance and Tishkoff 2013). We therefore decided to only retain these CHMM samples in the PCA and ADMIXTURE analyses, but to remove them from the subsequent analyses. When excluding the four samples from CHMM, we retrieved 30,034,755 variants, which after filtering, reduced to 16,383,002 biallelic SNPs and 1,179,117 LD-pruned biallelic SNPs. All of the samples from Thailand (*n* = 30) and Malaysia (*n* = 1), were found to be unrelated based on relatedness and kinship analyses (*Sup. Fig.S2*).

### Population structure and geneflow

When analysing all samples (*n*=35), the principal components analysis (PCA) showed distinct clustering of populations based on geographic origin (Fig. 1A) which had PC1 and PC2 loadings of 15.00% and 9.32%, respectively (Fig. 1B). Geographic structure was evident within Thailand and Malaysia, with clusters representing southernmost and mid-south Thailand, and the Kao Yai (KY) forest in the east of the country. The remaining cluster extended from the Western Forest Complex northwards (Fig. 1B). While these clusters closely reflect geographic origin, the distribution of samples in the PCA plot may indicate continuous genetic variation among regions, disrupted by gaps in sampling. Samples from CHMM were distinct from Thai and Malaysian samples and were inseparable from each other in the PCA, reflecting the observed ascertainment bias and consequent homogeneity among these sample genotypes.

**Figure 1.**
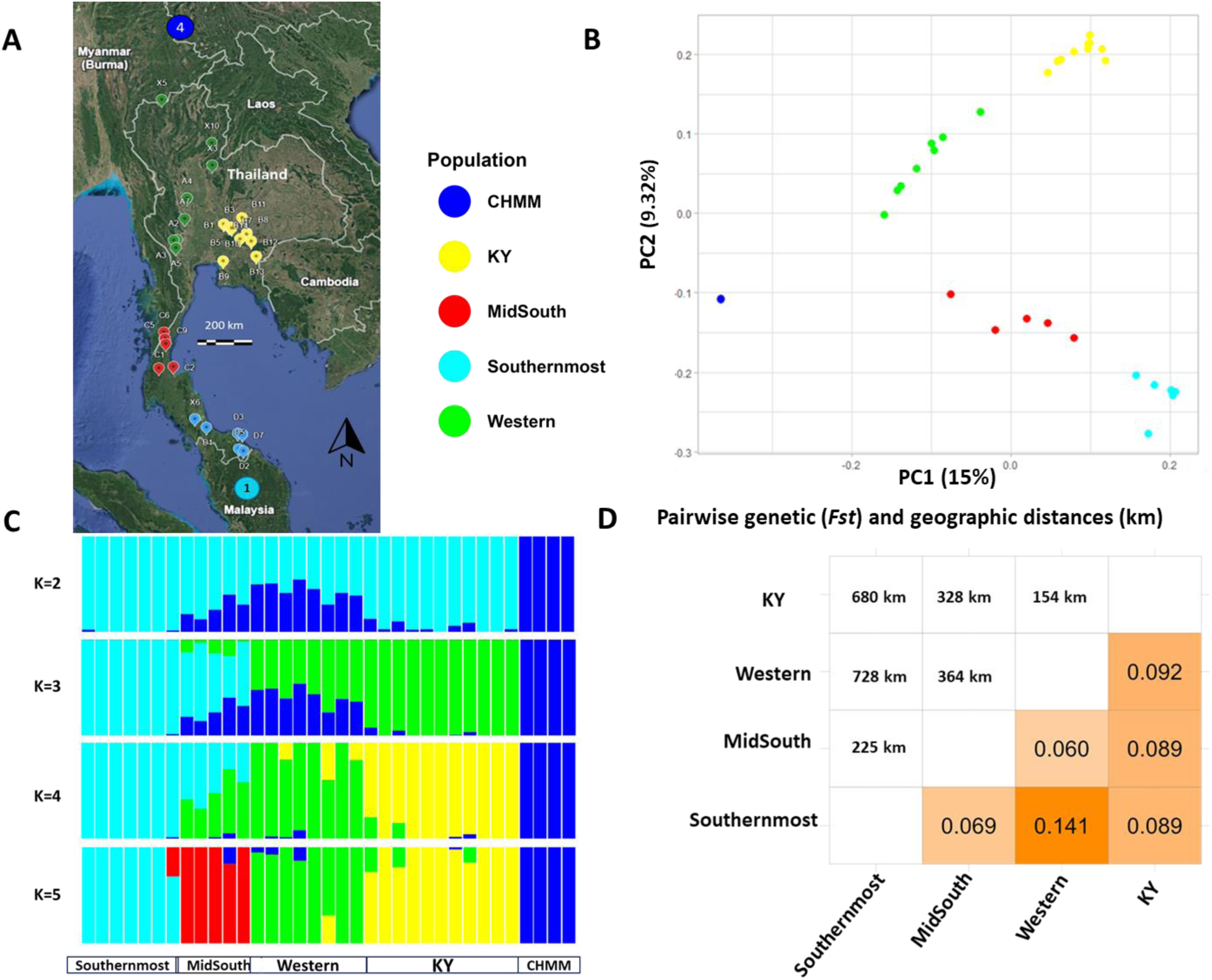
Sample locations and population structure of Sunda pangolin. ***A.*** Map of sample locations in this study (*n*=35): the smaller markers represent individual Sunda pangolins in Thailand, and their colours represent clusters according to PCA and ADMIXTURE analysis. The larger dark blue circle to the north represents four Sunda pangolins (three samples from Yunnan, China and one sample from Kachin, Myanmar), and the larger light blue circle to the south represents one Sunda pangolin from Malaysia. ***B***. Principal Component Analysis (PCA) for all 35 individuals distributed on the first two PC axes. *C*. Population structure inferred using ADMIXTURE analysis from *K* = 2 to *K* = 5. ***D.*** Genetic distance matrix based on *F*_ST_ among populations defined according to PCA and ADMIXTURE results (except CHMM), and geographic distances between each population (km), using the nearest individuals between each population pair.

The ADMIXTURE analysis indicated that the most likely number of genetic clusters is *K* = 2, based on the lowest cross-validation (CV) error (*Sup. Fig.S3*). At *K* = 2, there was clear differentiation between CHMM samples and the Thailand/Malaysia samples. At *K* = 5, the ADMIXTURE plot reflects the structure of five potential populations based on geographic locations in line with the PCA result (Fig. 1C): CHMM, Western Thailand (‘Western’), Central Thailand (Kao Yai Forest complex; ‘KY’), Mid-south Thailand (‘MidSouth’) and a ‘Southernmost’ population comprising samples from the far south of Thailand and one sample from the far north of Malaysia. Subsequent analyses focus on these four genetic populations observed in Thailand and Malaysia (*n = 31*), with the CHMM samples removed.

The *F_ST_* values between populations show moderate genetic differentiation, ranging from 0.06-0.14. In general, the scale of *F_ST_* corresponded to the geographic distance between each population (nearest sample localities); the lowest differentiation is found between Mid-South and Western populations (*F_ST_* = 0.060), while the Western and Southernmost population exhibited much higher differentiation (*F_ST_* = 0.141). However, the Western and Khao Yai (KY) populations, which are only approximately 154 km apart, showed a relatively high level of differentiation (*F_ST_* = 0.092) compared to the KY and Southernmost populations (*F_ST_* = 0.089) (Fig. 1D).

### Genetic diversity and inbreeding

There were no significant differences in nucleotide diversity (π) among populations (one-way ANOVA, p = 0.226) (Fig. 2A). The Southernmost population exhibited the highest mean nucleotide diversity (π = 0.000136), followed closely by the Mid-South (π = 0.000130) and KY (π = 0.000127) populations. The Western population displayed the lowest mean nucleotide diversity (π = 0.000119).

**Figure 2.**
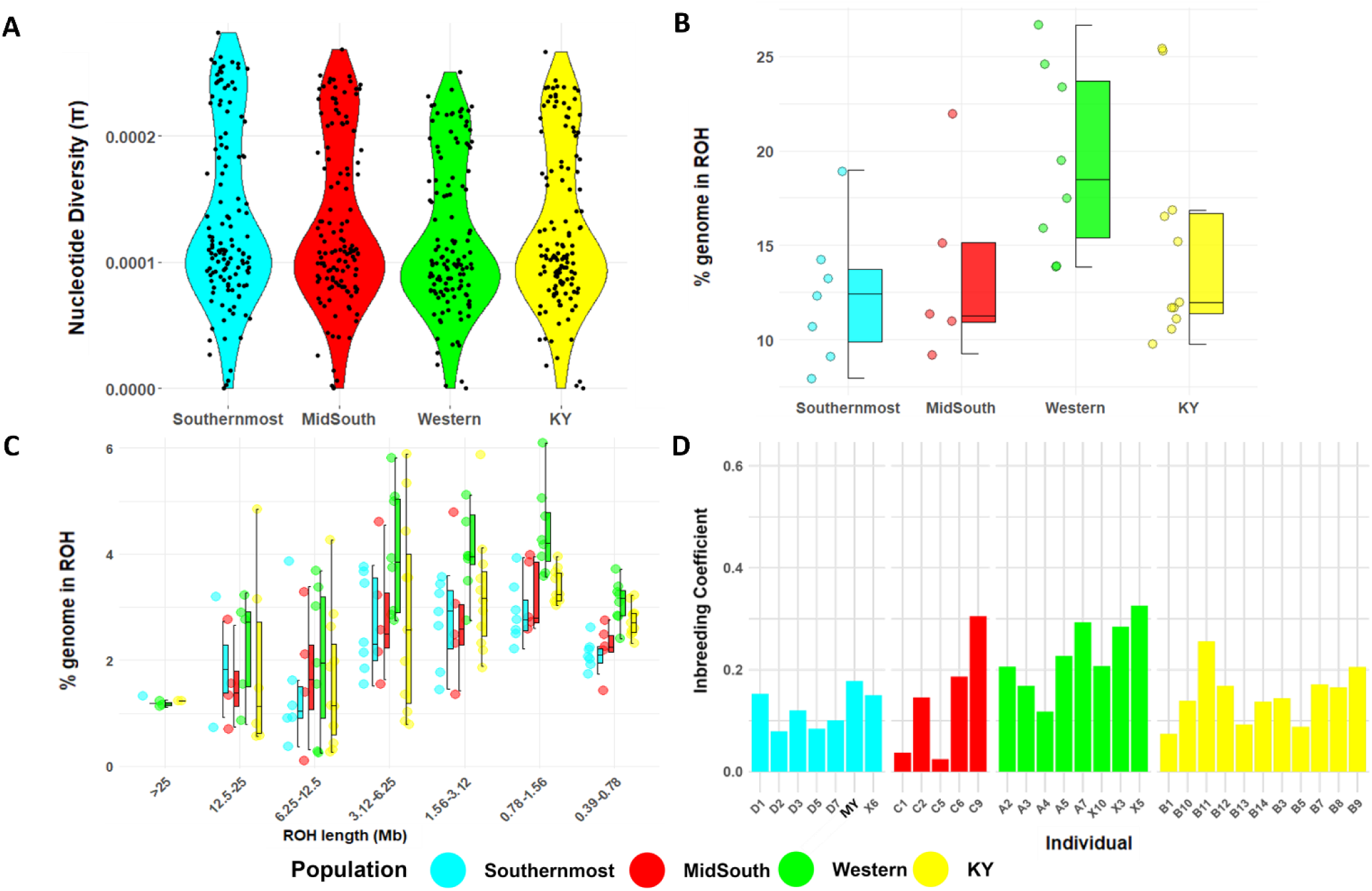
Genetic diversity and ROH distribution of Sunda pangolin. ***A.*** Nucleotide diversity in each population. Each data point represents the average nucleotide diversity across all samples at each of 20 Mb sliding windows. ***B***. Distribution of *F*_ROH_ across populations. The centre lines of boxplots reflect the median, the bounds of the boxes extend from the first to the third quartiles, and the upper and lower whiskers reflect the largest and smallest values but no further than 1.5 x the interquartile range from the hinge. The coloured dots on the left side of each boxplot reflect an individual in the population. ***C.*** Distribution of ROH within different length classes. Data points represent the percentage of ROH of a given length within an individual’s autosomal genome. ***D.*** Individual inbreeding coefficients in each population, calculated with VCFtools. Each individual label as the sample name exept one sample from Malaysia (SRR3949728) label as MY.

In contrast, significant differences were observed in heterozygosity among populations (one-way ANOVA, F3,27 = 11.47, p < 0.0001). Pairwise comparisons with Bonferroni correction showed that the Southernmost population (*He* = 0.00210 (± 0.000171)) had significantly higher heterozygosity than both Western (*He* = 0.00166 (± 0.000165); p <0.01) and KY populations (*He* = 0.00179 (± 0.000120); p = 0.001500), and the Western population had significantly lower heterozygosity than the MidSouth population (*He* = 0.00194 (±0.000160); p = 0.0229); other population pairs did not differ significantly (p > 0.05) (*Sup. Table S5*). A Shapiro-Wilk test confirmed that heterozygosity values were normally distributed in all populations (p > 0.05). A comparison of inter-species heterozygosity revealed that Sunda pangolin in Thailand has similar or slightly higher heterozygosity levels (*Sup. Fig.S4*) than the giant pangolin (*Smutsia gigantea*) (0.00187) and black-bellied pangolin (*Phataginus tetradactyla*) (0.00166) (Gu et al. 2023), however, these values remain notably lower than the white-bellied pangolin (*Phataginus tricuspis*) (0.00285) (*Sup. Fig.S2*), which exhibits the highest heterozygosity among pangolins.

Reduced diversity in the Western population was more evident from the ROH analysis, with the percentage of the genome in runs of homozygosity (*F_ROH_*) markedly higher in this population *F_ROH_* (18.50%) than the other three (all <12.5%; Fig. 2B). A Kruskal-Wallis test indicated that the differences in *F*_ROH_ were statistically significant among populations (χ^2^ = 7.869, df = 3, p = 0.0488), while, a Conover–Iman test revealed that this result was driven by the comparison between the Western population and the Southernmost population. Within populations, the Western population showed a wider range of *F*_ROH_ values among individuals, indicating more variability in the levels of homozygosity within this population. In contrast, the Southernmost population displays more consistent *F_ROH_* values across individuals.

The analysis of the ROH length class distribution across populations showed that the Western population exhibited higher median *F_ROH_* values in all classes, particularly when compared to the Southernmost population, which consistently showed the lowest *F*_ROH_ values (Fig. 2C). Individual inbreeding coefficients (*F_IS_*) (Fig. 2D) displayed considerable variation both within and between populations, with the four populations exhibiting a standard deviation ranging from 0.038 to 0.116. Consistent with *F*_ROH_, the Western population exhibited the highest mean *F_IS_* (0.229 ± 0.069), while the Southernmost population showed the lowest (0.124 ± 0.038). The MidSouth and KY populations displayed intermediate values (0.140 ± 0.116 and 0.149 ± 0.054, respectively) (*Sup. Table S6*). Additionally, the strong positive correlation between *F_ROH_* and *F_IS_* (Pearson’s r = 0.815, p < 0.000) further highlights the interplay between runs of homozygosity and inbreeding at the individual level (*Sup. Fig.S5*). AMOVA results showed that a significant level of total genetic variance was distributed among populations (*F* = 3.638, p = 0.025).

## Discussion

This study reveals genetic structure in the Sunda pangolin across Thailand, with four population genetic clusters corresponding to western, eastern, mid-south and southernmost sampling localities. Combined with the observed genetic variation among the populations, these results are set to inform pangolin conservation genetic management and the traceability of pangolin seizures within Thailand and neighbouring regions.

### Population structure

Our whole-genome SNP analysis reveals geographic population structure in Sunda pangolins that aligns with but also expands upon previous mitochondrial DNA (mtDNA) findings (Banterng et al. 2025), with greater resolution. The phylogeographic patterns observed in the previous mtDNA study showed five distinct clades in continental Southeast Asia with a clear north-south separation corresponding to the Kangar-Pattani Line, as well as an east-west split within Thailand between the western forest complex and Khao Yai forest complex populations. However, these mtDNA clades displayed introgression not observed in our nuclear genome data and in southern Thailand; the distribution of nuclear DNA-inferred populations and mtDNA-inferred clades is somewhat incongruent. The mid-south pangolins displayed clear nuclear genetic differentiation from central Thailand that was not observed in the mtDNA data, whereas in the southernmost region, the single population observed in the nuclear DNA analysis encompassed significant mtDNA lineage variation, suggesting differences between contemporary and historic evolutionary processes. Sunda pangolins from Yunnan Province (China) and Kachin (Myanmar) were distinct from other Southeast Asian samples, aligning with a previous study based on mtDNA haplotypes and nuclear SNPs (Hu et al. 2020).

The average pairwise genetic differentiation among populations (*F_ST_* = 0.09), indicates significant reductions in gene flow across the species range in Thailand. This level of differentiation is similar to that observed among adjacent populations in the Chinese pangolin (*F_ST_* = 0.104) occurring immediately north of the Sunda pangolin distribution (Wang et al. 2022). This suggests that historical and ecological factors common to this biogeographic region may have played a significant role in shaping population structure across multiple pangolin species.

Although the KY and Western populations are geographically close to each other (approximately 154 km apart), they showed moderate genetic differentiation (*F_ST_* = 0.093), suggesting a barrier to dispersal in central Thailand. In contrast to the discrete structure revealed by nuclear SNP markers, the previous mitogenome study showed some lineage introgression between these two regions (Banterng et al in press), suggesting that restrictions to gene flow between the Western Forest Complex and Kao Yai are likely relatively recent. The observed differentiation could be influenced by biogeographic and anthropogenic factors. The Chao Phraya River and associated major human settlements - including Bangkok and its surrounding agricultural communities - likely serve as barriers to dispersal. Historical events, such as the forced resettlement of over 200,000 prisoners of war approximately 200 years ago to relocate and build the new capital and increase its capacity for growing rice and harvesting timber for export (Baker and Phongpaichit 2022), may have further altered the landscape and connectivity between populations.

In southern Thailand, biogeographic barriers, such as the Nakhon Si Thammarat Mountain range, may also constitute important factors in shaping the genetic differentiation between the Mid-South and Southernmost populations (*F_ST_* = 0.069) despite their relatively close geographic proximity (*Sup. Fig.S6*). This mountain range is closely associated with the Kangar-Pattani Line (KPL), a significant biogeographical transition zone that demarcates the boundary between the Indochinese and Sundaic bioregions (Hughes, Round, and Woodruff 2003; Woodruff 2010). The overall genetic differentiation pattern strongly supports the role of the KPL as a barrier, though one samples appears to conflict with this pattern, suggesting that dispersal across this barrier is possible (*Sup. Fig.S6*). The patterns we have identified are similar to those observed in other mammalian species in the region, such as the grey-bellied (*Callosciurus caniceps*) and Asian red-cheeked (*Dremomys rufigenis*) squirrels (Hinckley et al. 2023), and white handed gibbons (*Hylobates lar*) (Gani et al. 2021). The genetic structure observed in our study in southern Thailand likely reflects historical separation due to geographical barriers, increased by recent anthropogenic pressures such as habitat loss and fragmentation.

### Genetic diversity

The observed differences in inbreeding levels and heterozygosity across populations highlight distinct genetic histories. The Western population exhibits the highest levels of inbreeding, with longer ROH and significantly lower heterozygosity compared to other populations, particularly the Southernmost population. This suggests a history of more recent inbreeding, potentially due to a smaller effective population size and/or restricted gene flow into the region. The gradual increase in heterozygosity from Western to Southernmost populations (Western < KY < MidSouth < Southernmost) aligns with inbreeding trends, suggesting a possible north-south gradient in genetic diversity.

Within the Western population, samples A7, X3, and X5 displayed the notably high levels of inbreeding (Fig. 2D) suggesting that these individuals may have experienced restricted mating opportunities due to specific local landscape features promoting limited dispersal (e.g., habitat fragmentation), or recent population declines in their immediate vicinity. In contrast, X10, the most geographically isolated member of the Western population, exhibits lower inbreeding. This unexpected pattern suggests that geographic distance alone may not determine genetic isolation, and potentially highlights limitations in our sampling, and/or variation in historic genetic diversity. Similarly, individual C9 from the MidSouth population exhibits high levels of inbreeding despite clustering geographically with other MidSouth individuals. This result emphasises the potential heterogeneity of genetic diversity within a population, which may be shaped by local environmental or historical events, such as localised population bottlenecks.

The Southernmost population exhibits the highest overall heterozygosity, the lowest inbreeding levels, and relatively uniform *F*_ROH_ values, indicative of a more genetically diverse and resilient population. This result is consistent with a previous study on the species’ mitogenome, which found that the southernmost region, a transboundary forest complex between Thailand and Malaysia, contains 17 haplotypes within this small area (Banterng et al in press), suggesting that this region may have maintained a consistently larger population over time, or been the recipient of gene flow from neighbouring populations, supporting its higher genetic diversity.

While overall nucleotide diversity remains relatively consistent across populations, differences in heterozygosity, homozygosity, and inbreeding levels reveal key genetic patterns. These findings emphasise the importance of considering heterozygosity and inbreeding alongside nucleotide diversity when assessing population genetic health. Even populations with similar nucleotide diversity can differ significantly in their levels of heterozygosity and inbreeding history, with important implications for long-term genetic viability.

A previous study that examined Sunda pangolin genetic diversity identified two distinct genetic lineages, which were putatively assigned to the north of the species’ continental distribution (China, Myanmar; designated ‘MJA’) and their distribution further south on the Southeast Asian peninsula and neighbouring islands (designated ‘MJB’), including Thailand (Hu et al. 2020). The *F_ROH_* and SNP heterozygosity estimates of the MJB lineage are broadly similar to the measurements in our study; the MJA lineage exhibited much lower levels of diversity. The slight discrepancy in heterozygosity between MJB and the pangolins in our study may be due to differences in individual sampling within this lineage, or could potentially stem from methodological variations. We used genome-wide approaches to account for SNP uncertainty and incorporated both polymorphic and monomorphic sites, instead of estimating heterozygosity based only on SNP sites (called using VCFtools) (Hu et al. 2020). Indeed, SNP-based heterozygosity estimates can be biased by sample size and population structure, whereas genome-wide approaches that include all sites (invariable and variant sites) provide robust and standardised estimates (Schmidt et al. 2021).

Using genome-wide SNP heterozygosity allowed us to robustly compare diversity across different species. Our results indicate that Sunda pangolin genetic diversity is comparable to that of the giant and black-bellied pangolins, while the Philippines pangolin (*Manis culionensis*) was lower and the white-bellied pangolin considerably higher (*Sup. Fig.S2*). The Sunda pangolin exhibited heterozygosity levels most similar to endangered primate species such as the Western gorilla and the chimpanzee from Central Africa. Despite all pangolin species being either Critically Endangered or Endangered due to their rapid, recent human-induced declines, their genome-wide heterozygosity values are not particularly low when compared with other mammals (*Sup. Fig.S2*), corresponding to an expected lag in the reduction of heterozygosity compared to demography.

### Implications for conservation units and traceability

Our fine-scale analysis of population structure and genetic diversity across the Sunda pangolin range in Thailand underscores the value of genomic approaches in providing important information for developing effective conservation strategies that should maintain genetic diversity and ensure the long-term viability of endangered populations (Coates, Byrne, and Moritz 2018; Moritz 2002). Interpretation should be considered within the context of our limited sample size (*n = 30*); expanded sampling would further elucidate whether the patterns observed represent discrete populations or reflect continuous variation with isolation by distance effects. Nevertheless, the genetic differentiation observed between populations in Thailand, coupled with varying levels of inbreeding and genetic diversity suggest that these populations should be considered as distinct Management Units (MUs) and can be applied to wildlife conservation and management strategies (Hohenlohe, Funk, and Rajora 2021). For instance, the Western population, with its higher inbreeding levels and lower heterozygosity, may benefit from genetic rescue programs to increase diversity (Ralls et al. 2020), while the more genetically diverse Southernmost population should be prioritised for habitat protection to maintain its genetic variability (Moritz 2002). Additionally, the identification of geographical barriers like the Nakhon Si Thammarat Mountain range and the Chao Phraya River as factors influencing genetic structure, could be used to inform policymakers of effective conservation strategies for Sunda pangolins in the future.

Notably, our study identified genetic populations show strong alignment with Thailand’s established forest complex system managed by the Department of National Parks, Wildlife and Plant Conservation (*Sup. Fig.S7*) (Emphandhu and Chettamart 2003). Correspondence between the MUs characterised here and administrative management units provides a valuable framework for implementing conservation strategies. Specifically, the Western population predominantly occurs within the Western Forest complex with one individual in the Lum Nam Pai-Salawin forest complex and one in the Phu Meang–Phu Thong Forest Complex, while the MidSouth population is primarily distributed across the Chumphon and Klong Saaeng-Khao Sok forest complexes. The Southernmost population, which showed higher genetic diversity, is mainly found within the Khao Bantad and Hala-Bala Forest complexes, and the KY population primarily inhabits the Dong Phayayen-Khao Yai Forest complex. The significant overlap between genetic populations and existing forest complex boundaries indicates these administrative units could effectively serve as management units for conservation. This alignment enables the integration of genetic-based conservation strategies within the established management framework.

Our finding of distinct population structure also has significant implications for developing traceability tools and enhancing wildlife forensic capabilities. The genetic differences between populations provide a promising foundation for creating a SNP panel that could serve as a powerful forensic tool in combating illegal wildlife trade (Ogden 2011). Similar SNP-based traceability approaches have proven successful in other pangolin species, with Nash et al. (2018) demonstrating how >12,000 SNPs could effectively assign seized Sunda pangolins to their geographic origins across insular Southeast Asia, revealing previously unrecognised genetic lineages and trade patterns. More recently, Tinsman et al. (2023) developed a panel of 96 diagnostic SNPs that successfully traced white-bellied pangolin seizures to specific geographic origins with high precision (median error of 72.2 km).

## Conclusion

Our findings have significant implications for the conservation and management of the Sunda pangolin, providing novel insights into their population structure and genetic diversity. This study serves as a foundational baseline for understanding the genetic landscape of this Critically Endangered species in Thailand, and addressing a significant knowledge gap in Sunda pangolin biogeography. The genetic information uncovered here can inform targeted conservation strategies, guide management decisions, and contribute to the development of more effective protection measures. Moreover, the population structure can be leveraged to develop traceability tools for wildlife forensics, aiding in the fight against illegal trafficking of this species. Given the limited information previously available on this species, our results offer a valuable contribution to the scientific understanding of Sunda pangolins and provide a solid basis for future conservation efforts.

## Data availability

The sequencing data are deposited in the NCBI under the BioProject accession

PRJNA1240953: Genetic analysis and traceability of Sunda pangolins in Thailand; SRR32831660 to SRR32831689.

## Acknowledgement

We would like to thank Emily Humble, Melissa Marr, Simone D’Alessandro and Marc-Alexander Gose from University of Edinburgh, for the bioinformatic consultation. Xier Luo and colleagues from Foshan University, for provided the pseudo-chromosome reference genome, Prateep Duengkae from Kasetsart University for advice. We also thank all staff at DNP Thailand who helped collect the samples in this project.

## Author Contributions

NB performed laboratory work and data analysis. RO conceived the study, supervised lab work and data analysis. NB, RO and KE wrote the manuscript. All authors read and approved the manuscript.

## Corresponding author

Correspondence to Nattapong Banterng and Rob Ogden

## References

Alexander, David H, John Novembre, and Kenneth Lange. 2009. ‘Fast model-based estimation of ancestry in unrelated individuals’, Genome research, 19: 1655–64.

Allendorf, Fred W., Paul A. Hohenlohe, and Gordon Luikart. 2010. ‘Genomics and the future of conservation genetics’, Nature Reviews Genetics, 11: 697–709.

Baker, Chris, and Pasuk Phongpaichit. 2022. ‘The old order in transition, 1760s to 1860s.’ in Chris Baker and Pasuk Phongpaichit (eds.), A History of Thailand (Cambridge University Press: Cambridge).

Banterng, et al, Kyle. Ewart, Frankie Sitam, Rob Ogden. in press. ‘Crossing boundaries: Mitogenomic analysis of Thai Sunda pangolins reveals regional phylogeography, and informs conservation management’, Scientific Reports.

Banterng, Nattapong, Kyle Ewart, Frankie Thomas Sitam, and Rob Ogden. 2025. ‘Mitogenomic analysis of Thai Sunda pangolins reveals regional phylogeography and informs conservation management’, Scientific Reports, 15: 14067.

Barbosa, Soraia, Sarah A. Hendricks, W. Chris Funk, Om P. Rajora, and Paul A. Hohenlohe. 2021. ’Wildlife Population Genomics: Applications and Approaches.’ in Paul A. Hohenlohe and Om P. Rajora (eds.), Population Genomics: Wildlife (Springer International Publishing: Cham).

Challender, D, DHA Willcox, E Panjang, N Lim, H Nash, S Heinrich, and J Chong. 2019. ‘Manis javanica. The IUCN red list of threatened species 2019: e. T12763A123584856." In.

Challender, Daniel W. S., Sarah Heinrich, Chris R. Shepherd, and Lydia K. D. Katsis. 2020. ‘Chapter 16 - International trade and trafficking in pangolins, 1900–2019.’ in Daniel W. S. Challender, Helen C. Nash and Carly Waterman (eds.), Pangolins (Academic Press).

Chong, Ju Lian, Elisa Panjang, Daniel Willcox, Helen C. Nash, Gono Semiadi, Withoon Sodsai, Norman T. L. Lim, Louise Fletcher, Ade Kurniawan, and Shavez Cheema. 2020. ‘Chapter 6 - Sunda pangolin Manis javanica (Desmarest, 1822).’ in Daniel W. S. Challender, Helen C. Nash and Carly Waterman (eds.), Pangolins (Academic Press).

Coates, David J., Margaret Byrne, and Craig Moritz. 2018. ‘Genetic Diversity and Conservation Units: Dealing With the Species-Population Continuum in the Age of Genomics’, Frontiers in ecology and evolution, 6.

Danecek, Petr, Adam Auton, Goncalo Abecasis, Cornelis A. Albers, Eric Banks, Mark A. DePristo, Robert E. Handsaker, Gerton Lunter, Gabor T. Marth, Stephen T. Sherry, Gilean McVean, Richard Durbin, and Genomes Project Analysis Group. 2011. ‘The variant call format and VCFtools’, Bioinformatics, 27: 2156–58.

Danecek, Petr, James K Bonfield, Jennifer Liddle, John Marshall, Valeriu Ohan, Martin O Pollard, Andrew Whitwham, Thomas Keane, Shane A McCarthy, Robert M Davies, and Heng Li. 2021. ‘Twelve years of SAMtools and BCFtools’, GigaScience, 10.

Emphandhu, Dachanee, and Surachet Chettamart. 2003. ‘Thailand’s experience in protected area management’, Faculty of Forestry, Kasetsart University, Bangkok, Thailand.

Ewart, Kyle M., Amanda L. Lightson, Frankie T. Sitam, Jeffrine Rovie-Ryan, Son G. Nguyen, Kelly I. Morgan, Adrian Luczon, Edwin Miguel S. Anadon, Marli De Bruyn, Stéphanie Bourgeois, Kanita Ouitavon, Antoinette Kotze, Mohd Soffian A. Bakar, Milena Salgado-Lynn, and Ross McEwing. 2021. ‘DNA analyses of large pangolin scale seizures: Species identification validation and case studies’, Forensic Science International: Animals and Environments, 1: 100014.

Flanagan, Sarah P., Brenna R. Forester, Emily K. Latch, Sally N. Aitken, and Sean Hoban. 2018. ‘Guidelines for planning genomic assessment and monitoring of locally adaptive variation to inform species conservation’, Evolutionary Applications, 11: 1035–52.

Forester, Brenna R., Erin L. Landguth, Brian K. Hand, and Niko Balkenhol. 2021. ’Landscape Genomics for Wildlife Research.’ in Paul A. Hohenlohe and Om P. Rajora (eds.), Population Genomics: Wildlife (Springer International Publishing: Cham).

Funk, W. Chris, John K. McKay, Paul A. Hohenlohe, and Fred W. Allendorf. 2012. ‘Harnessing genomics for delineating conservation units’, Trends in Ecology & Evolution, 27: 489–96.

Gani, Millawati, Jeffrine J Rovie-Ryan, Frankie Thomas Sitam, Noor Azleen Mohd Kulaimi, Chew Cheah Zheng, Aida Nur Atiqah, Nur Maisarah Abd Rahim, and Ahmad Azhar Mohammed. 2021. ‘Taxonomic and genetic assessment of captive White-Handed Gibbons (Hylobateslar) in Peninsular Malaysia with implications towards conservation translocation and reintroduction programmes’, ZooKeys, 1076: 25.

Gu, Tong-Tong, Hong Wu, Feng Yang, Philippe Gaubert, Sean P. Heighton, Yeyizhou Fu, Ke Liu, Shu-Jin Luo, Hua-Rong Zhang, Jing-Yang Hu, and Li Yu. 2023. ‘Genomic analysis reveals a cryptic pangolin species’, Proceedings of the National Academy of Sciences, 120: e2304096120.

Gu, Tongtong, Jingyang Hu, and Li Yu. 2024. ‘Evolution and conservation genetics of pangolins’, Integrative Zoology, 19: 426–41.

Hanghøj, Kristian, Ida Moltke, Philip Alstrup Andersen, Andrea Manica, and Thorfinn Sand Korneliussen. 2019. ‘Fast and accurate relatedness estimation from high-throughput sequencing data in the presence of inbreeding’, GigaScience, 8.

Heighton, Sean P, Rémi Allio, Jérôme Murienne, Jordi Salmona, Hao Meng, Céline Scornavacca, Armanda DS Bastos, Flobert Njiokou, Darren W Pietersen, and Marie-Ka Tilak. 2023. ‘Pangolin genomes offer key insights and resources for the world’s most trafficked wild mammals’, Molecular biology and evolution, 40: msad190.

Heighton, Sean P., and Philippe Gaubert. 2021. ‘A timely systematic review on pangolin research, commercialization, and popularization to identify knowledge gaps and produce conservation guidelines’, Biological Conservation, 256: 109042.

Hinckley, Arlo, Melissa TR Hawkins, Jesús E Maldonado, and Jennifer A Leonard. 2023. ‘Evolutionary history and patterns of divergence in three tropical east Asian squirrels across the Isthmus of Kra’, Journal of Biogeography, 50: 1090–102.

Hohenlohe, Paul A., W. Chris Funk, and Om P. Rajora. 2021. ‘Population genomics for wildlife conservation and management’, Molecular ecology, 30: 62–82.

Hu, Jing-Yang, Zi-Qian Hao, Laurent Frantz, Shi-Fang Wu, Wu Chen, Yun-Fang Jiang, Hong Wu, Wei-Min Kuang, Haipeng Li, Ya-Ping Zhang, and Li Yu. 2020. ‘Genomic consequences of population decline in critically endangered pangolins and their demographic histories’, National Science Review, 7: 798–814.

Hughes, Jennifer B., Philip D. Round, and David S. Woodruff. 2003. ‘The Indochinese–Sundaic faunal transition at the Isthmus of Kra: an analysis of resident forest bird species distributions’, Journal of Biogeography, 30: 569–80.

Humble, Emily, Martin A. Stoffel, Kara Dicks, Alex D. Ball, Rebecca M. Gooley, Justin Chuven, Ricardo Pusey, Mohammed Al Remeithi, Klaus-Peter Koepfli, Budhan Pukazhenthi, Helen Senn, and Rob Ogden. 2023. ‘Conservation management strategy impacts inbreeding and mutation load in scimitar-horned oryx’, Proceedings of the National Academy of Sciences, 120: e2210756120.

Islam, Rabiul, Yefang Li, Xuexue Liu, Haile Berihulay, Adam Abied, Gebremedhin Gebreselassie, Qing Ma, and Yuehui Ma. 2019. ‘Genome-Wide Runs of Homozygosity, Effective Population Size, and Detection of Positive Selection Signatures in Six Chinese Goat Breeds’, Genes, 10: 938.

Korneliussen, Thorfinn Sand, Anders Albrechtsen, and Rasmus Nielsen. 2014. ‘ANGSD: Analysis of Next Generation Sequencing Data’, BMC Bioinformatics, 15: 356.

Lachance, Joseph, and Sarah A. Tishkoff. 2013. ‘SNP ascertainment bias in population genetic analyses: Why it is important, and how to correct it’, BioEssays, 35: 780–86.

Li, Heng, and Richard Durbin. 2009. ‘Fast and accurate short read alignment with Burrows–Wheeler transform’, Bioinformatics, 25: 1754–60.

Li, Heng, and Richard Durbin. 2011. ‘Inference of human population history from individual whole-genome sequences’, Nature, 475: 493–96.

Marth, Gabor T, Eva Czabarka, Janos Murvai, and Stephen T Sherry. 2004. ‘The Allele Frequency Spectrum in Genome-Wide Human Variation Data Reveals Signals of Differential Demographic History in Three Large World Populations’, Genetics, 166: 351–72.

Moritz, Craig. 2002. ‘Strategies to Protect Biological Diversity and the Evolutionary Processes That Sustain It’, Systematic biology, 51: 238–54.

Nash, Helen C., Wirdateti Wirdateti, Gabriel W. Low, Siew Woh Choo, Ju Lian Chong, Gono Semiadi, Ranjeev Hari, Muhammad Hafiz Sulaiman, Samuel T. Turvey, Theodore A. Evans, and Frank E. Rheindt. 2018. ‘Conservation genomics reveals possible illegal trade routes and admixture across pangolin lineages in Southeast Asia’, Conservation genetics, 19: 1083–95.

Nielsen, Rasmus, Melissa J Hubisz, and Andrew G Clark. 2004. ‘Reconstituting the Frequency Spectrum of Ascertained Single-Nucleotide Polymorphism Data’, Genetics, 168: 2373–82.

Ogden, Rob. 2011. ‘Unlocking the potential of genomic technologies for wildlife forensics’, Molecular Ecology Resources, 11: 109–16.

Ogden, Rob, and Adrian Linacre. 2015. ‘Wildlife forensic science: A review of genetic geographic origin assignment’, Forensic Science International: Genetics, 18: 152–59.

Poplin, Ryan, Pi-Chuan Chang, David Alexander, Scott Schwartz, Thomas Colthurst, Alexander Ku, Dan Newburger, Jojo Dijamco, Nam Nguyen, and Pegah T Afshar. 2018. ‘A universal SNP and small-indel variant caller using deep neural networks’, Nature biotechnology, 36: 983–87.

Presti, Flavia T., Neiva M. R. Guedes, Paulo T. Z. Antas, and Cristina Y. Miyaki. 2015. ‘Population Genetic Structure in Hyacinth Macaws (Anodorhynchus hyacinthinus) and Identification of the Probable Origin of Confiscated Individuals’, Journal of Heredity, 106: 491–502.

Priyambada, Prajnashree, Gul Jabin, Abhishek Singh, Avijit Ghosh, Sujeet Kumar Singh, Moitrye Chatterjee, Chinnadurai Venkatraman, Kailash Chandra, Lalit Kumar Sharma, and Mukesh Thakur. 2021. ‘Digging out the keys in the heap of seized pangolin scales: up scaling pangolin conservation using wildlife forensics’, Forensic Science International, 323: 110780.

Purcell, Shaun, Benjamin Neale, Kathe Todd-Brown, Lori Thomas, Manuel AR Ferreira, David Bender, Julian Maller, Pamela Sklar, Paul IW De Bakker, and Mark J Daly. 2007. ‘PLINK: a tool set for whole-genome association and population-based linkage analyses’, The American journal of human genetics, 81: 559–75.

Ralls, Katherine, Paul Sunnucks, Robert C. Lacy, and Richard Frankham. 2020. ‘Genetic rescue: A critique of the evidence supports maximizing genetic diversity rather than minimizing the introduction of putatively harmful genetic variation’, Biological Conservation, 251: 108784.

Ritland, Kermit. 1996. ‘Estimators for pairwise relatedness and individual inbreeding coefficients’, Genetics Research, 67: 175–85.

Santiago, Enrique, Irene Novo, Antonio F Pardiñas, María Saura, Jinliang Wang, and Armando Caballero. 2020. ‘Recent Demographic History Inferred by High-Resolution Analysis of Linkage Disequilibrium’, Molecular biology and evolution, 37: 3642–53.

Schmidt, Thomas L, Moshe-Elijah Jasper, Andrew R Weeks, and Ary A Hoffmann. 2021. ‘Unbiased population heterozygosity estimates from genome-wide sequence data’, Methods in Ecology and Evolution, 12: 1888–98.

Shirley, Matthew H., Georgina Gerard, Elisa Panjang, Nick Ching-Min Sun, and Sean P. Heighton. 2023. ‘Pangolins: epitomizing the complexities of conservation’, Oryx, 57: 681–82.

Sitam, Frankie T., Milena Salgado-Lynn, Azroie Denel, Elisa Panjang, Ross McEwing, Amanda Lightson, Rob Ogden, Nur Alwanie Maruji, Nurhartini Kamalia Yahya, Cosmas Ngau, Noor Azleen Mohd Kulaimi, Hartini Ithnin, Jeffrine Rovie-Ryan, Mohd Soffian Abu Bakar, and Kyle M. Ewart. 2023. ‘Phylogeography of the Sunda pangolin, Manis javanica: Implications for taxonomy, conservation management and wildlife forensics’, Ecology and Evolution, 13: e10373.

Theissinger, Kathrin, Carlos Fernandes, Giulio Formenti, Iliana Bista, Paul R. Berg, Christoph Bleidorn, Aureliano Bombarely, Angelica Crottini, Guido R. Gallo, José A. Godoy, Sissel Jentoft, Joanna Malukiewicz, Alice Mouton, Rebekah A. Oomen, Sadye Paez, Per J. Palsbøll, Christophe Pampoulie, María J. Ruiz-López, Simona Secomandi, Hannes Svardal, Constantina Theofanopoulou, Jan de Vries, Ann-Marie Waldvogel, Guojie Zhang, Erich D. Jarvis, Miklós Bálint, Claudio Ciofi, Robert M. Waterhouse, Camila J. Mazzoni, Jacob Höglund, Sargis A. Aghayan, Tyler S. Alioto, Isabel Almudi, Nadir Alvarez, Paulo C. Alves, Isabel R. Amorim do Rosario, Agostinho Antunes, Paula Arribas, Petr Baldrian, Giorgio Bertorelle, Astrid Böhne, Andrea Bonisoli-Alquati, Ljudevit L. Boštjančić, Bastien Boussau, Catherine M. Breton, Elena Buzan, Paula F. Campos, Carlos Carreras, L. FIlipe C. Castro, Luis J. Chueca, Fedor Čiampor, Elena Conti, Robert Cook-Deegan, Daniel Croll, Mónica V. Cunha, Frédéric Delsuc, Alice B. Dennis, Dimitar Dimitrov, Rui Faria, Adrien Favre, Olivier D. Fedrigo, Rosa Fernández, Gentile Francesco Ficetola, Jean-François Flot, Toni Gabaldón, Dolores R. Agius, Alice M. Giani, M. Thomas P. Gilbert, Tine Grebenc, Katerina Guschanski, Romain Guyot, Bernhard Hausdorf, Oliver Hawlitschek, Peter D. Heintzman, Berthold Heinze, Michael Hiller, Martin Husemann, Alessio Iannucci, Iker Irisarri, Kjetill S. Jakobsen, Peter Klinga, Agnieszka Kloch, Claudius F. Kratochwil, Henrik Kusche, Kara K. S. Layton, Jennifer A. Leonard, Emmanuelle Lerat, Gianni Liti, Tereza Manousaki, Tomas Marques-Bonet, Pável Matos-Maraví, Michael Matschiner, Florian Maumus, Ann M. Mc Cartney, Shai Meiri, José Melo-Ferreira, Ximo Mengual, Michael T. Monaghan, Matteo Montagna, Robert W. Mysłajek, Marco T. Neiber, Violaine Nicolas, Marta Novo, Petar Ozretić, Ferran Palero, Lucian Pârvulescu, Marta Pascual, Octávio S. Paulo, Martina Pavlek, Cinta Pegueroles, Loïc Pellissier, Graziano Pesole, Craig R. Primmer, Ana Riesgo, Lukas Rüber, Diego Rubolini, Daniele Salvi, Ole Seehausen, Matthias Seidel, Bruno Studer, Spyros Theodoridis, Marco Thines, Lara Urban, Anti Vasemägi, Adriana Vella, Noel Vella, Sonja C. Vernes, Cristiano Vernesi, David R. Vieites, Christopher W. Wheat, Gert Wörheide, Yannick Wurm, and Gabrielle Zammit. 2023. ‘How genomics can help biodiversity conservation’, Trends in Genetics, 39: 545–59.

Thompson, Elizabeth A. 2013. ‘Identity by descent: variation in meiosis, across genomes, and in populations’, Genetics, 194: 301–26.

Tinsman, Jen C., Cristian Gruppi, Christen M. Bossu, Tracey-Leigh Prigge, Ryan J. Harrigan, Virginia Zaunbrecher, Klaus-Peter Koepfli, Matthew LeBreton, Kevin Njabo, Cheng Wenda, Shuang Xing, Katharine Abernethy, Gary Ades, Excellence Akeredolu, Imuzei B. Andrew, Taneisha A. Barrett, Iva Bernáthová, Barbora Černá Bolfíková, Joseph L. Diffo, Ghislain Difouo Fopa, Lionel Esong Ebong, Ichu Godwill, Aurélie Flore Koumba Pambo, Kim Labuschagne, Julius Nwobegahay Mbekem, Brice R. Momboua, Carla L. Mousset Moumbolou, Stephan Ntie, Elizabeth Rose-Jeffreys, Franklin T. Simo, Keerthana Sundar, Markéta Swiacká, Jean Michel Takuo, Valery N. K. Talla, Ubald Tamoufe, Caroline Dingle, Kristen Ruegg, Timothy C. Bonebrake, and Thomas B. Smith. 2023. ‘Genomic analyses reveal poaching hotspots and illegal trade in pangolins from Africa to Asia’, Science, 382: 1282–86.

Wang, Qing, Tianming Lan, Haimeng Li, Sunil Kumar Sahu, Minhui Shi, Yixin Zhu, Lei Han, Shangchen Yang, Qian Li, Le Zhang, Zhangwen Deng, Huan Liu, and Yan Hua. 2022. ‘Whole-genome resequencing of Chinese pangolins reveals a population structure and provides insights into their conservation’, Communications Biology, 5: 821.

Waples, Robin S., and Eric C. Anderson. 2017. ‘Purging putative siblings from population genetic data sets: a cautionary view’, Molecular ecology, 26: 1211–24.

Wasser, Samuel K., Charles J. Wolock, Mary K. Kuhner, John E. Brown, Chris Morris, Ryan J. Horwitz, Anna Wong, Charlene J. Fernandez, Moses Y. Otiende, Yves Hoareau, Zofia A. Kaliszewska, Eunjin Jeon, Kin-Lan Han, and Bruce S. Weir. 2022. ‘Elephant genotypes reveal the size and connectivity of transnational ivory traffickers’, Nature Human Behaviour, 6: 371–82.

Wong, Rebecca W. Y. 2019. The Illegal Wildlife Trade in China Understanding The Distribution Networks / by Rebecca W. Y. Wong (Springer International Publishing: Cham).

Woodruff, David S. 2010. ‘Biogeography and conservation in Southeast Asia: how 2.7 million years of repeated environmental fluctuations affect today’s patterns and the future of the remaining refugial-phase biodiversity’, Biodiversity and Conservation, 19: 919–41.

Yan, Dingyu, Xier Luo, Jiabin Tang, Shanghua Xu, Kongwei Huang, Xiaobo Wang, Tong Feng, Tengcheng Que, Miaomiao Jia, Xiaobing Guo, Saif ur Rehman, Zhipeng Li, Yufeng Yang, Kaixiang Li, Kuiqing Cui, Jue Ruan, and Qingyou Liu. 2022. ‘High-Quality Genomes of Pangolins: Insights into the Molecular Basis of Scale Formation and Adaption to Myrmecophagous Diet’, Molecular biology and evolution, 40.

Yun, Taedong, Helen Li, Pi-Chuan Chang, Michael F Lin, Andrew Carroll, and Cory Y McLean. 2021. ‘Accurate, scalable cohort variant calls using DeepVariant and GLnexus’, Bioinformatics, 36: 5582–89.

Zhang, H. 2020. ‘Genetic identification of African pangolins and their origin in illegal trade’, Global Ecol. Conserv., 23: e01119.

Zhang, Huarong, Gary Ades, Mark P. Miller, Feng Yang, Kwok-wai Lai, and Gunter A. Fischer. 2020. ‘Genetic identification of African pangolins and their origin in illegal trade’, Global ecology and conservation, 23: e01119.

Zhang, Huarong, Mark P. Miller, Feng Yang, Hon Ki Chan, Philippe Gaubert, Gary Ades, and Gunter A. Fischer. 2015. ‘Molecular tracing of confiscated pangolin scales for conservation and illegal trade monitoring in Southeast Asia’, Global ecology and conservation, 4: 414–22.

