## Supplementary Materials for "Genomic Population Structure of the Sunda Pangolin (*Manis javanica*) in Thailand: Implications for Conservation and Wildlife Forensics"

**Table S1.** Details of Sunda pangolins sequenced in this study, along with Accession numbers of sequence data obtained. The samples obtained in the current study derived from the Department of National Parks, Wildlife and Plant Conservation (DNP), Thailand.

| Sample name | NCBI accession No. | Geographic provenance | Collection | Sampling locality | Sample type | Study | Source |
|---|---|---|---|---|---|---|---|
| A2 | SRR32831689 | Western forest complex | DNP | Salakpra Wildlife Sanctuary | Blood | Current study | Rescued |
| A3 | SRR32831688 | Western forest complex | DNP | Salakpra Wildlife Sanctuary | Blood | Current study | Rescued |
| A4 | SRR32831677 | Western forest complex | DNP | Ladyao, Nakhon Sawan | Blood | Current study | Rescued |
| A5 | SRR32831666 | Western forest complex | DNP | Khaoson Wildlife breeding station | Tissue | Current study | Rescued |
| A7 | SRR32831665 | Western forest complex | DNP | Krasiao Dam, Suphan Buri | Blood | Current study | Rescued |
| B1 | SRR32831664 | Khao Yai forest complex | DNP | Saraburi | Blood | Current study | Rescued |
| B3 | SRR32831663 | Khao Yai forest complex | DNP | Tublan National Park | Blood | Current study | Rescued |
| B5 | SRR32831662 | Khao Yai forest complex | DNP | Panggae, Pakchong | Blood | Current study | Rescued |
| B7 | SRR32831661 | Khao Yai forest complex | DNP | Na Di, Prachin Buri | Blood | Current study | Rescued |
| B8 | SRR32831660 | Khao Yai forest complex | DNP | Tublan National Park | Blood | Current study | Rescued |
| B9 | SRR32831687 | Khao Yai forest complex | DNP | Nong Prue, Bang Lamung | Blood | Current study | Rescued |
| B10 | SRR32831686 | Khao Yai forest complex | DNP | Khao Yai National Park | Blood | Current study | Rescued |

| | | | | | | | |
|---|---|---|---|---|---|---|---|
| B11 | SRR32831685 | Khao Yai forest complex | DNP | Si Khio,Nakhon Ratchasima | Blood | Current study | Rescued |
| B12 | SRR32831684 | Khao Yai forest complex | DNP | Khok Pee Khong, Sa Kaeo | Blood | Current study | Rescued |
| B13 | SRR32831683 | Khao Yai forest complex | DNP | Klonghad, Sa Kaeo | Blood | Current study | Rescued |
| B14 | SRR32831682 | Khao Yai forest complex | DNP | Krabin Buri, Prachin Buri | Blood | Current study | Rescued |
| C1 | SRR32831681 | Mid-south, Thailand | DNP | Khirirat Nikhom, Surat Thani | Blood | Current study | Rescued |
| C2 | SRR32831680 | Mid-south, Thailand | DNP | Makham Tia,Surat Thani | Blood | Current study | Rescued |
| C5 | SRR32831679 | Mid-south, Thailand | DNP | Thamsing, Chumphon | Blood | Current study | Rescued |
| C6 | SRR32831678 | Mid-south, Thailand | DNP | Ban Na, Chumphon | Tissue | Current study | Rescued |
| C9 | SRR32831676 | Mid-south, Thailand | DNP | Langsuan, Chumphon | Tissue (Hair) | Current study | Rescued |
| D1 | SRR32831675 | Southernmost, Thailand | DNP | Ton Nga Chang Wildlife Sanctuary | Blood | Current study | Rescued |
| D2 | SRR32831674 | Southernmost, Thailand | DNP | Buketa, Narathiwat | Tissue | Current study | Rescued |
| D3 | SRR32831673 | Southernmost, Thailand | DNP | Khok Khian, Narathiwat | Tissue | Current study | Rescued |
| D5 | SRR32831672 | Southernmost, Thailand | DNP | Hala Bala Wildlife Sanctuary | Tissue | Current study | Confiscated |
| D7 | SRR32831671 | Southernmost, Thailand | DNP | Khok Khian, Narathiwat | Blood | Current study | Rescued |

| | | | | | | | |
|---|---|---|---|---|---|---|---|
| X6 | SRR32831668 | Southernmost, Thailand | DNP | Palian, Trang | Blood | Current study | Rescued |
| X3 | SRR32831670 | Central, Thailand | DNP | Kaeng Sopha, Phitsanulok | Blood | Current study | Rescued |
| X5 | SRR32831669 | Northern, Thailand | DNP | Chiang Dao, Wildlife Sanctuary | Blood | Current study | Rescued |
| X10 | SRR32831667 | Northern, Thailand | DNP | Huai Phueng Wang Yao Non-Hunting Area | Blood | Current study | Rescued |
| SRR9018633 | SRR9018633 | - | GenBank | Kachin, Myanmar | - | Hu et al., 2020 | - |
| SRR9018664 | SRR9018664 | - | GenBank | Yunnan Province, China | - | Hu et al., 2020 | - |
| SRR9018665 | SRR9018665 | - | GenBank | Yunnan Province, China | - | Hu et al., 2020 | - |
| SRR9018658 | SRR9018658 | - | GenBank | Yunnan Province, China | - | Hu et al., 2020 | |
| SRR3949728 | SRR3949728 | - | GenBank | Malaysia | - | Choo et al. 2016 | |

**Table S2**. Data used to compare genome-wide heterozygosity, modified from Marr et al. 2025.

| **Species** | **Common name** | **Heterozygosity** | **Status** | **Coverage** | **Source** | **DOI** |
|---|---|---|---|---|---|---|
| *Panthera tigris altaica* | Amur tiger | 0.00061 | EN | high > 20x | Cho et al. 2013 | 4:2433doi:10.1038/ncomms 3433(2013). |
| *Lipotes vexillifer* | Baiji | 0.000121 | CR | high > 20x | Zhou et al. 2013 | 10.1038/ncomms3708 |
| *Ovis canadensis* | Bighorn sheep | 0.002217 | LC | high > 20x | Corbett-Detig et al. 2015 | 10.1371/journal.pbio.10021 12 |
| *Phataginus tetradactyla* | Black-bellied pangolin (PTE20) | 0.001664 | VU | high > 20x | Gu et al. 2024 | https://doi.org/10.1073/pna s.2304096120 |
| *Pheobastria nigripes* | Black-footed albatross | 0.0008 | NT | low (5x) | Huynh et al. 2023 | https://doi.org/10.1093/mol bev/msad155 |
| *Acinonyx jubatus* | Cheetah | 0.0002 | VU | low 5 - 6x | Dobrynin et al. 2015 | 10.1186/s13059-015-0837-4 |
| *Pan troglodytes troglodytes* | Chimpanzee (Central African) | 0.00176 | EN | high > 20x | The Chimpanzee Sequencing and Analysis Consortium 2005 | 10.1038/nature04072 |
| *Pan troglodytes verus* | Chimpanzee (West African) | 0.00087 | EN | high > 20x | The Chimpanzee Sequencing and Analysis Consortium 2005 | 10.1038/nature04072 |
| *Sciurus vulgaris* | Eurasian red squirrel | 0.0002 | LC | low 3.7 -7.5x | Marr et al. 2025 | https://doi.org/10.1111/eva. 70072 |
| *Balaenoptera physalus* | Fin whale | 0.00151 | VU | high > 20x | Yim et al. 2014 | 10.1038/ng.2835 |
| *Ailuropoda melanoleuca* | Giant panda | 0.00135 | VU | high > 20 | Ruiqiang Li et al. 2010 | https://doi.org/10.1038/nat ure08696 |
| *Smutsia gigantea* | Giant pangolin (SGI20) | 0.001879 | EN | low - mid (9x) | Gu et al. 2023 | https://doi.org/10.1073/pna s.2304096120 |
| *Lynx pardinus* | Iberian Lynx | 0.00022 | EN | high > 20x | Abascal et al. 2016 | https://doi.org/10.1186/s13 059-016-1090-1 |

| | | | | | | |
|---|---|---|---|---|---|---|
| *Manis crassicaudata* | Indian pangolin (MCR10) | 0.001011 | EN | high > 20x | Gu et al. 2023 | https://doi.org/10.1073/pnas.2304096120 |
| *Neofelis nebulosa* | Mainland clouded leopard | 0.000406 | VU | high > 20x | Yuan et al. 2023 | 10.1126/sciadv.adh9143 |
| *Gorilla beringei beringei* | Mountain gorilla | 0.000647 | CR | high > 20x | Xue et al. 2015 | 10.1126/science.aaa3952 |
| *Ovibos moschatus* | Musk-ox (Greenland) | 0.000125 | LC | low (4x) | Hansen et al. 2018 | 10.1016/j.cub.2018.10.054 |
| *Oreamnos americanus* | North American Mountain Goat | 0.000897 | LC | low - mid (9x) | Martchenko et al. 2023 | https://doi.org/10.1038/s41437-023-00643-4 |
| *Manis culionensis* | Philippine pangolin (MCU) | 0.000401 | CR | low 2x | Hu et al. 2020 | https://doi.org/10.1093/nsr/nwaa031 |
| *Equus ferus przewalskii* | Przewalski's horse | 0.0036 | EN | high > 20x | Corbett-Detig et al. 2015 | 10.1371/journal.pbio.1002112 |
| *Panthera uncia* | Snow leopard | 0.00023 | EN | high > 20x | Cho et al. 2013 | doi:10.1038/ncomms3433(2013). |
| *Pongo abelii* | Sumatran orangutan | 0.0012 | CR | high > 20x | Cho et al. 2013 | doi:10.1038/ncomms3433(2013). |
| *Manis javanica* | Sunda pangolin (A5) | 0.001706 | CR | low - mid (9x) | This study | |
| *Manis javanica* | Sunda pangolin (B12) | 0.001799 | CR | low - mid (9x) | This study | |
| *Manis javanica* | Sunda pangolin (C2) | 0.001925 | CR | low - mid (9x) | This study | |
| *Manis javanica* | Sunda pangolin (D3) | 0.002102 | CR | low - mid (9x) | This study | |
| *Sarcophilus harrisii* | Tasmanian devil | 0.00032 | EN | high > 20x | Miller et al. 2011 | 10.1073/pnas.1102838108 |
| *Smutsia temminckii* | Temminck 's pangolin (STE02) | 0.000974 | VU | high > 20x | Gu et al. 2023 | https://doi.org/10.1073/pnas.2304096120 |
| *Gorilla gorilla* | Western gorilla | 0.0019 | CR | high > 20x | Xue et al. 2015 | 10.1126/science.aaa3952 |

| | | | | | | |
|---|---|---|---|---|---|---|
| *Gorilla gorilla gorilla* | Western lowland gorilla | 0.001438 | CR | high > 20x | Xue et al. 2015 | 10.1126/science.aaa3952 |
| *Panthera leo* | White African lion | 0.00053 | VU | high > 20x | Armstrong et al. 2020 | https://doi.org/10.1186/s12915-019-0734-5 |
| *Phataginus tricuspis* | White-bellied pangolin (PTR09) | 0.002857 | EN | high > 20x | Gu et al. 2023 | https://doi.org/10.1073/pnas.2304096120 |
| *Gulo gulo* | Wolverine | 0.00028 | VU | low - mid 8-11x | Ekblom et al. 2018 | https://doi.org/10.1111/cobi.13157 |
| *Pan paniscus* | Bonobo | 0.0008 | EN | high > 20x | Prado-Martinez et al. 2013 | https://doi.org/10.1038/nature12228 |

**Table S3.** Variant calling and sequencing depth summary, result from VCF file using DeepVariant v 1.5.0. (Ti/Tv ratio = Transition/Transversion ratio).

| Sample | Number of sites | Average depth | Biallelic SNP | Ti/Tv ratio | Reference calls |
|---|---|---|---|---|---|
| A2 | 13944256 | 11.90 | 8415064 | 2.19 | 4459734 |
| A3 | 13878904 | 12.27 | 8732672 | 2.18 | 4031678 |
| A4 | 14772486 | 14.24 | 8908191 | 2.18 | 4712172 |
| A5 | 14899631 | 12.58 | 8411230 | 2.19 | 5420170 |
| A7 | 12573012 | 10.84 | 8017723 | 2.19 | 3544053 |
| B1 | 14171641 | 13.68 | 8771476 | 2.19 | 4264155 |
| B3 | 12626269 | 11.09 | 8209240 | 2.19 | 3386631 |
| B5 | 14305108 | 12.95 | 8574322 | 2.19 | 4641948 |
| B7 | 13390735 | 11.40 | 8200480 | 2.19 | 4159111 |
| B8 | 13904530 | 19.15 | 8667787 | 2.17 | 4081596 |
| B9 | 12424206 | 11.09 | 7975562 | 2.19 | 3442277 |
| B10 | 12829826 | 11.70 | 8330735 | 2.18 | 3444577 |
| B11 | 12836213 | 11.41 | 7946455 | 2.19 | 3893345 |
| B12 | 12254874 | 10.33 | 7973739 | 2.18 | 3288035 |
| B13 | 13618520 | 13.30 | 8540424 | 2.19 | 3978030 |
| B14 | 12415803 | 10.80 | 8129613 | 2.19 | 3262394 |
| C1 | 14551136 | 15.01 | 9116534 | 2.19 | 4249979 |
| C2 | 12279360 | 10.13 | 8222881 | 2.19 | 3024438 |
| C5 | 14572567 | 15.93 | 9211499 | 2.19 | 4163937 |
| C6 | 14190030 | 11.62 | 8340388 | 2.20 | 4802038 |
| C9 | 12677165 | 9.79 | 7611698 | 2.20 | 4131966 |
| D1 | 12827141 | 12.89 | 8434318 | 2.19 | 3309243 |
| D2 | 15490470 | 14.46 | 8744327 | 2.20 | 5628118 |
| D3 | 13750249 | 11.85 | 8287990 | 2.20 | 4426013 |
| D5 | 14415931 | 13.15 | 8582755 | 2.20 | 4750031 |
| D7 | 12367318 | 10.86 | 8332235 | 2.19 | 2988706 |
| SRR3949728 | 12258185 | 15.40 | 9388796 | 2.13 | 1662360 |
| SRR9018633 | 12341756 | 15.48 | 8125282 | 2.00 | 3115210 |
| SRR9018658 | 10311457 | 16.46 | 7895818 | 2.15 | 1417043 |
| SRR9018664 | 11260337 | 16.52 | 7961305 | 2.15 | 2273388 |
| SRR9018665 | 11117498 | 16.86 | 7919582 | 2.15 | 2166839 |
| X3 | 11570977 | 9.99 | 7921955 | 2.18 | 2653251 |
| X5 | 12035833 | 10.45 | 7953131 | 2.17 | 3082105 |
| X6 | 12866927 | 11.34 | 8160586 | 2.20 | 3685921 |
| X10 | 13264148 | 12.25 | 8471966 | 2.18 | 3712610 |

**Table S4.** Samples and sequencing data from BAM files derived from BWA-MEM alignment against the reference genome (GCA_024605085.1). The genome-wide heterozygosity estimates derived from ANGSD analysis with quality filtering (Phred Q20, proper pairs only) and controlled read depth (1-21×) using folded site frequency spectrum.

| Sample | Clean data (bp) | Clean data depth | Mapping (%) | Mapping depth | Heterozygosity |
|---|---|---|---|---|---|
| A2 | 23514264366 | 9.63 | 98.35 | 9.68933 | 0.001797 |
| A3 | 24220748057 | 9.92 | 98.46 | 9.96961 | 0.001802 |
| A4 | 29172787111 | 11.95 | 98.49 | 11.9718 | 0.001894 |
| A5 | 26774714676 | 10.97 | 98.11 | 10.9335 | 0.001706 |
| A7 | 20549359075 | 8.42 | 98.86 | 8.50679 | 0.001527 |
| B1 | 29184087392 | 11.96 | 98.57 | 11.9962 | 0.001901 |
| B3 | 21023056615 | 8.61 | 98.99 | 8.69735 | 0.001887 |
| B5 | 26034240005 | 10.66 | 98.8 | 10.6861 | 0.001977 |
| B7 | 22359075601 | 9.16 | 98.46 | 9.21198 | 0.001847 |
| B8 | 40176527895 | 16.46 | 98.95 | 16.5091 | 0.001653 |
| B9 | 22025408861 | 9.02 | 99.15 | 9.10484 | 0.001641 |
| B10 | 22431273093 | 9.19 | 99.02 | 9.26783 | 0.001855 |
| B11 | 21750005995 | 8.91 | 98.93 | 8.99104 | 0.001595 |
| B12 | 19161102812 | 7.85 | 99.11 | 7.94835 | 0.001799 |
| B13 | 27847728779 | 11.41 | 99.08 | 11.4795 | 0.001796 |
| B14 | 21515501507 | 8.81 | 99.22 | 8.88673 | 0.001759 |
| C1 | 30856047938 | 12.64 | 98.89 | 12.6938 | 0.002095 |
| C2 | 19939532929 | 8.17 | 98.65 | 8.25137 | 0.001925 |
| C5 | 32909780181 | 13.48 | 99.14 | 13.5355 | 0.002102 |
| C6 | 23211777509 | 9.51 | 98.76 | 9.31817 | 0.001809 |
| C9 | 18147693314 | 7.43 | 99.02 | 7.52554 | 0.001753 |
| D1 | 26365698227 | 10.8 | 99.05 | 10.8765 | 0.001843 |
| D2 | 29605116265 | 12.13 | 98.43 | 12.0925 | 0.002229 |
| D3 | 23401817389 | 9.59 | 98.34 | 9.66029 | 0.002102 |
| D5 | 26073326782 | 10.68 | 98.59 | 10.7094 | 0.002191 |
| D7 | 21529399789 | 8.82 | 99.27 | 8.8936 | 0.001983 |
| SRR3949728 | 33376280462 | 13.67 | 98.15 | 13.8377 | 0.002346 |
| X10 | 24088872441 | 9.87 | 99.05 | 9.93321 | 0.001666 |
| X3 | 19603949220 | 8.03 | 99.13 | 8.12171 | 0.001464 |
| X5 | 19539737581 | 8 | 99.04 | 8.08794 | 0.001463 |
| X6 | 21635286568 | 8.86 | 98.91 | 8.94692 | 0.001996 |

**Table S5.** Genome-wide heterozygosity by population.

| Population | Samples | Mean heterozygosity | Standard deviation | Minimum heterozygosity | Maximum heterozygosity |
|---|---|---|---|---|---|
| KY | 11 | 0.00179 | 0.000120 | 0.00160 | 0.00198 |
| MidSouth | 5 | 0.00194 | 0.000160 | 0.00175 | 0.00210 |
| Southernmost | 7 | 0.00210 | 0.000171 | 0.00184 | 0.00235 |
| Western | 8 | 0.00166 | 0.000165 | 0.00146 | 0.00189 |

**Note**: The distribution of heterozygosity values was normal for all populations (Shapiro-Wilk test, $p > 0.05$). One-way ANOVA revealed significant differences in heterozygosity among populations ($F3,27 = 11.47$, $p < 0.0001$).

**Table S6.** Summary statistics of inbreeding coefficients across four populations of Sunda pangolin in this study.

| Population | Samples | Mean Inbreeding Coefficient ($F_{IS}$) | Standard Deviation | Range |
|---|---|---|---|---|
| Southernmost | 7 | 0.124 | 0.038 | 0.079-0.178 |
| MidSouth | 5 | 0.14 | 0.116 | 0.024-0.305 |
| Western | 8 | 0.229 | 0.069 | 0.118-0.325 |
| KY | 11 | 0.149 | 0.054 | 0.074-0.256 |

**Note**: Overall, mean inbreeding coefficient across all populations was 0.163 ± 0.076 (SD). Within-population variance (0.0046) was greater than between-population variance (0.0017). One-way ANOVA revealed significant differences among populations ($F = 3.638$, df = 3, 27, $p = 0.0252$).

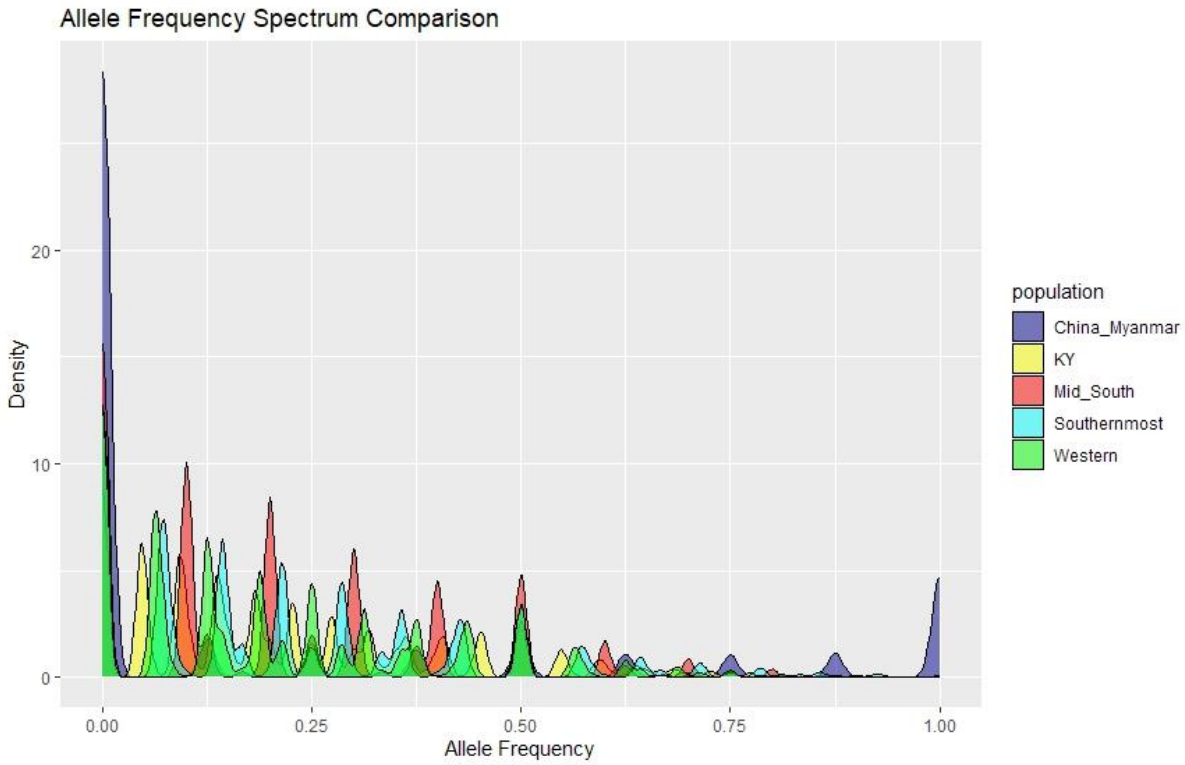


**Figure S1**. Allele frequency spectrum comparison among five populations in this study revealing ascertainment bias for China and Myanmar population.

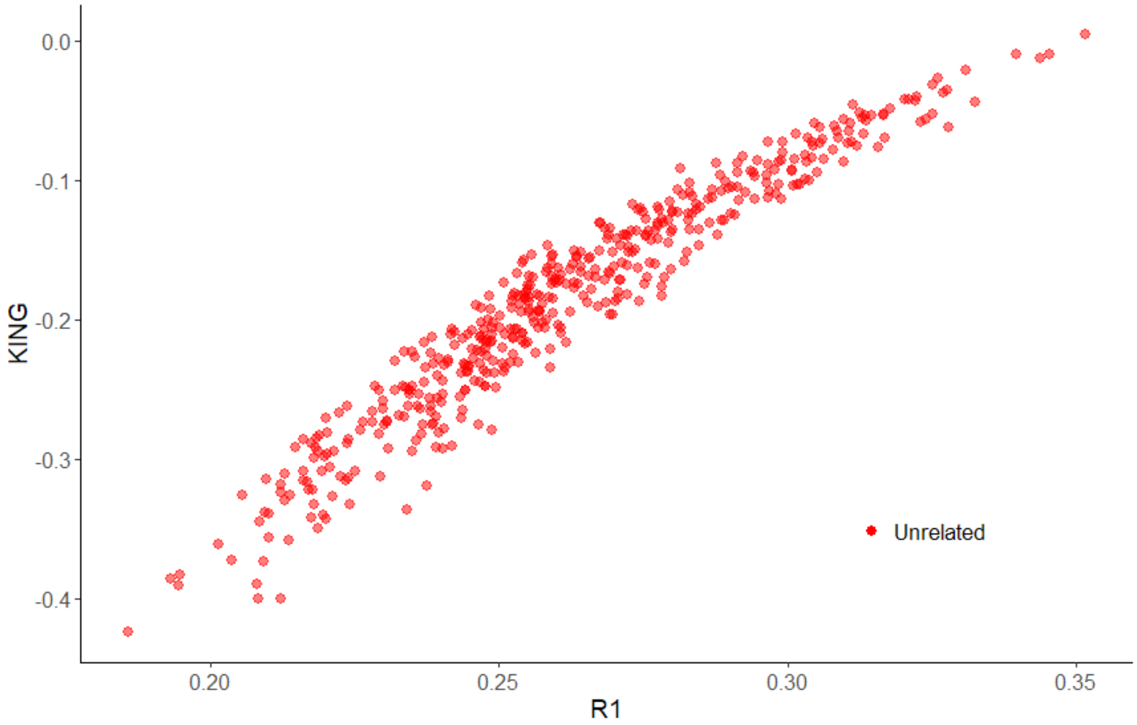


**Figure S2**. KING robust relatedness against R1 of pairwise comparisons between individual samples. Classification of relatedness category based on thresholds defined by Manichaikul et al. (2010), indicates that the 30 samples from Thailand and one sample from Malaysia in this study are not close-relatives.

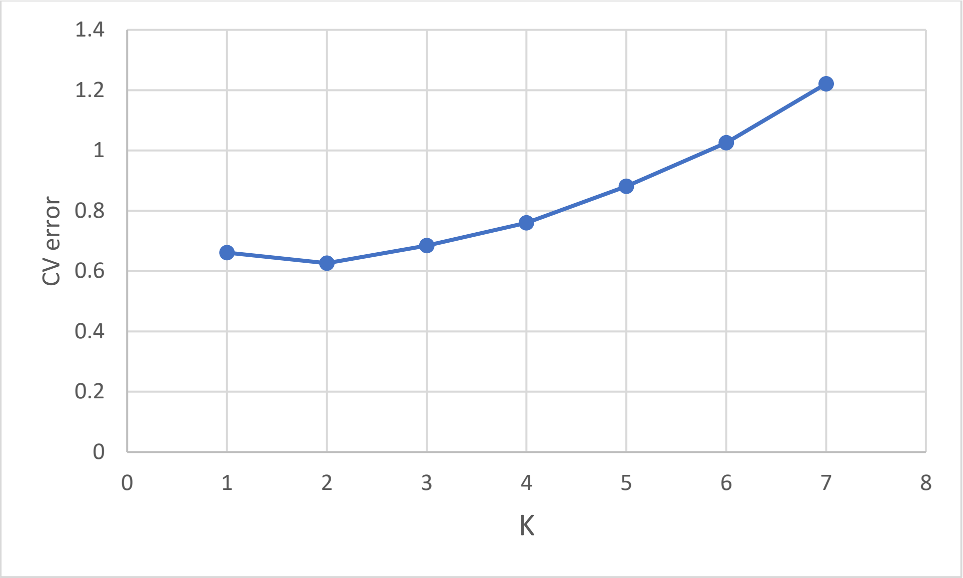


**Figure S3.** The cross-validation (CV) error from ADMIXTURE analysis, *K*=1-7, indicating the most likely number of genetic clusters in this study of Thai Sunda pangolins is *K*=2.

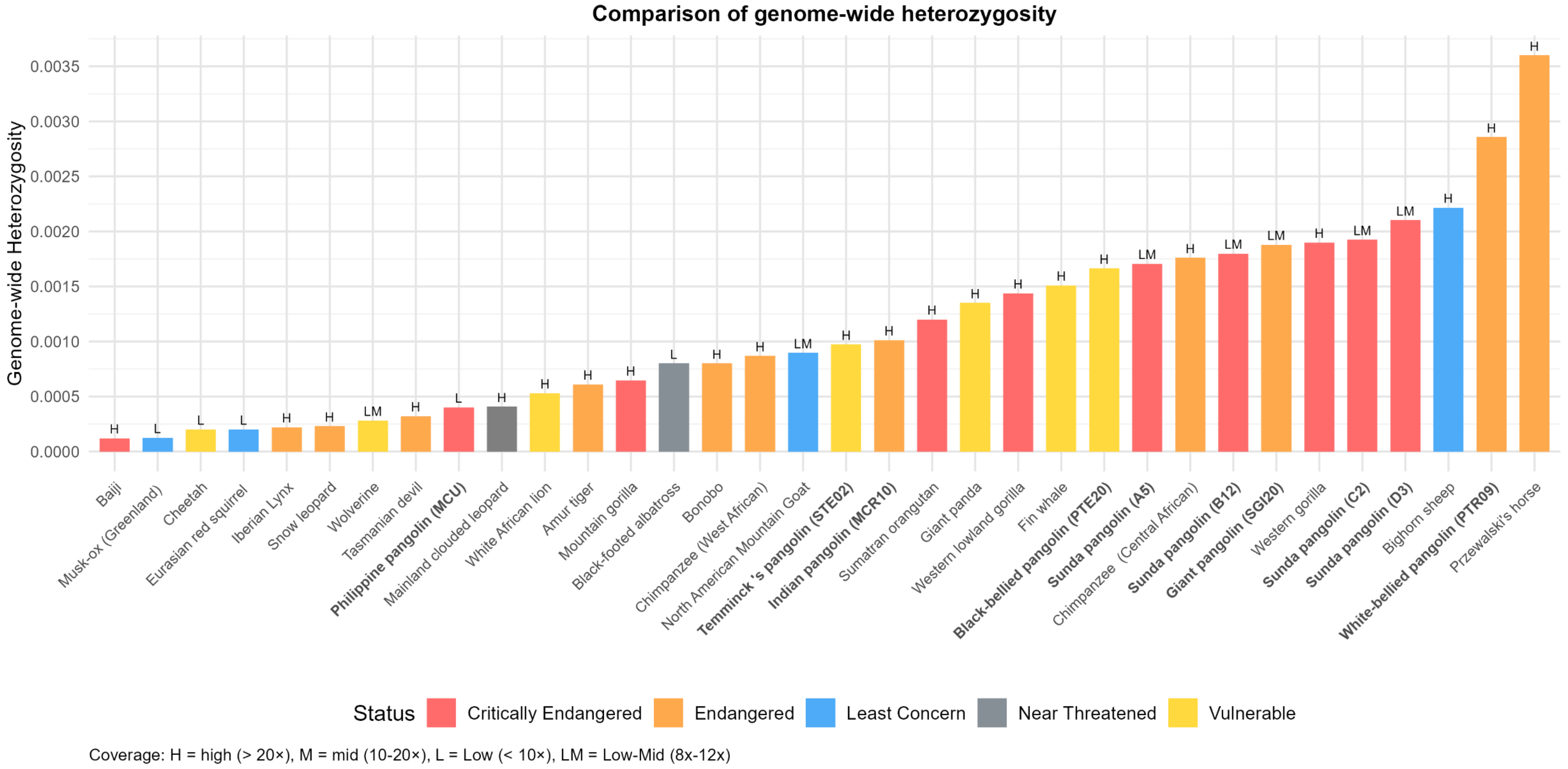


**Figure S4.** Comparison of genome-wide heterozygosity of pangolin species with other mammal and bird species with IUCN Status that represent in different colour. Genome-wide X-fold coverage: H = high (> 20×), M = mid (10–20×), L = Low (< 10×). Data modified from Marr et al. (2025) and references therein (Table S2). Note that these comparisons are approximations due to differences in datasets and variant calling.

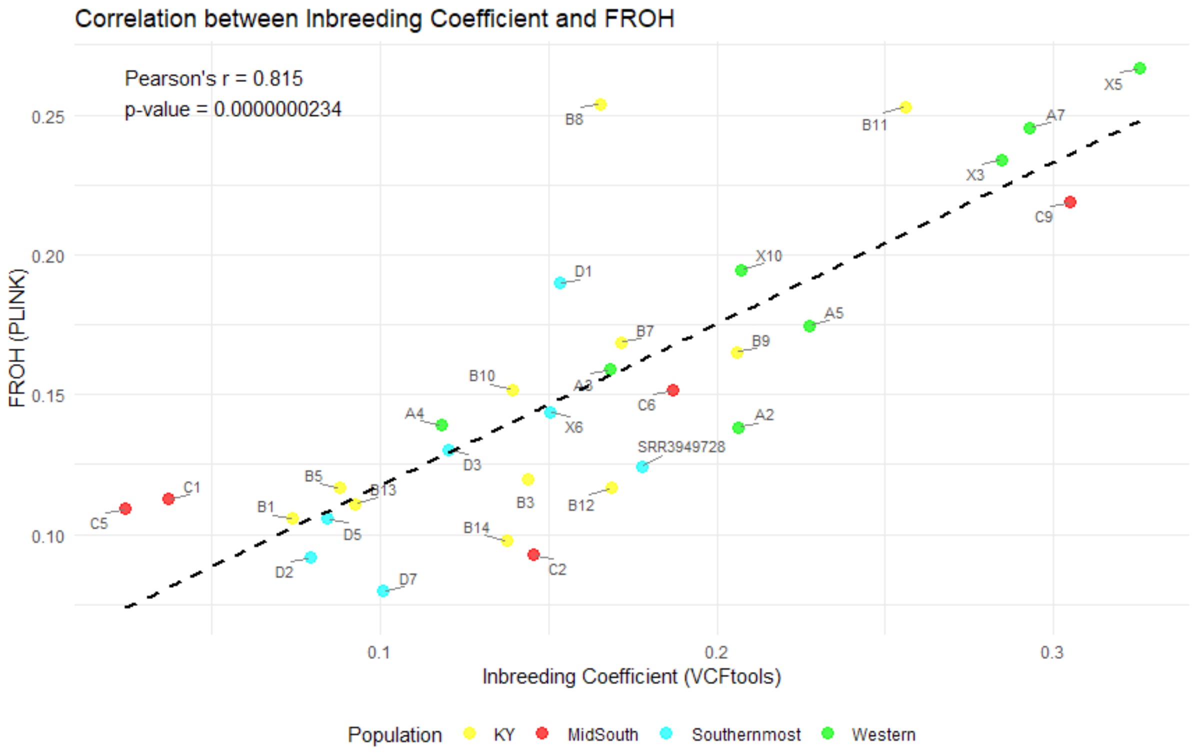


**Figure S5.** The correlation between $F_{ROH}$ from PLINK and inbreeding coefficient from VCFtools, Pearson's r = 0.815, p = 0.0000000234.

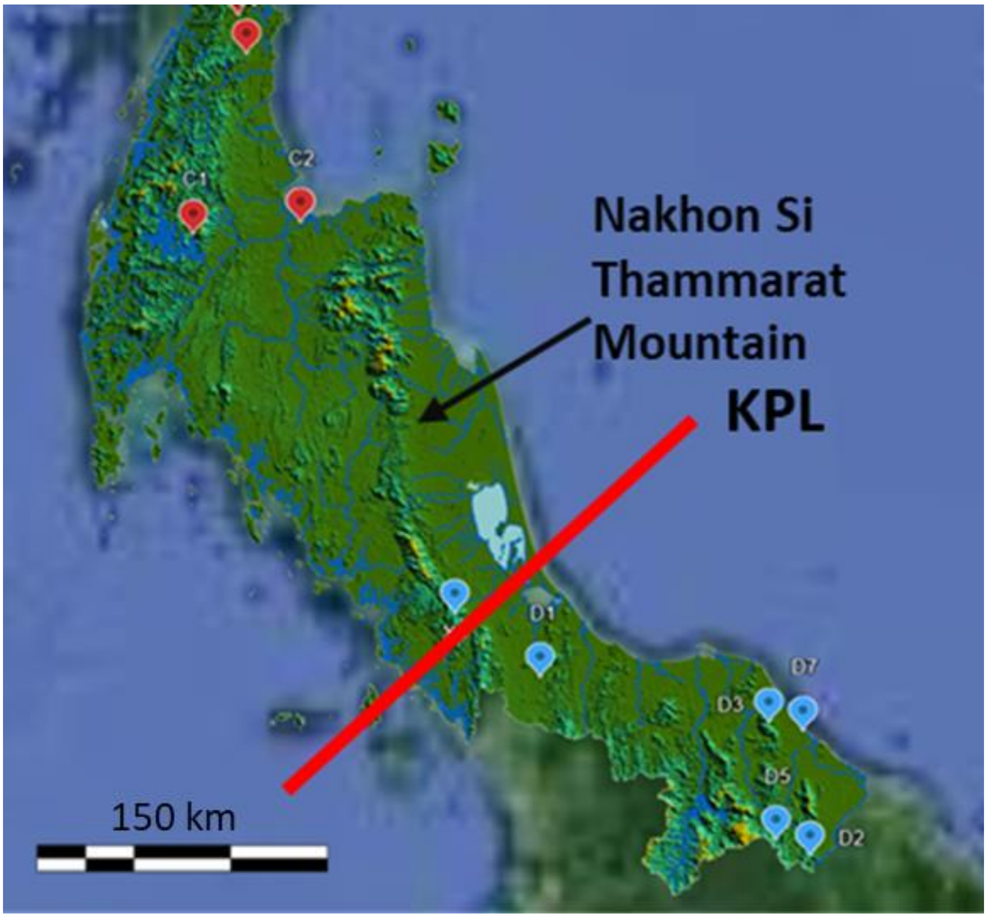


**Figure S6.** Map of the south of Thailand showing the location of the Nakhon Si Thammarat Mountain, the Kangar Pattani Line and the individual locations for the mid-south (red mark) and the southernmost (blue mark) pangolins.

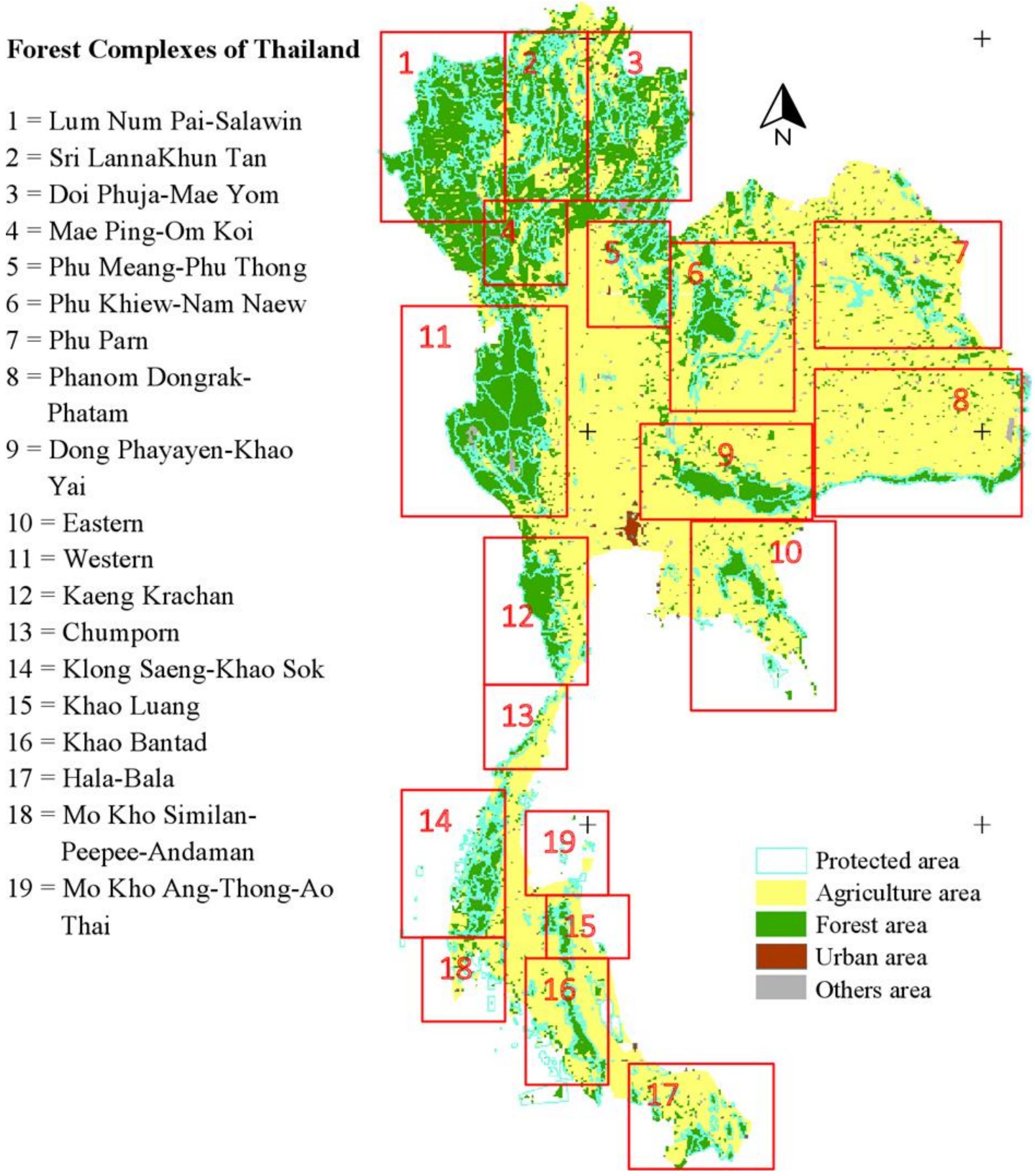


**Figure S7.** The forest complexes of Thailand (red squares), according to the Department of National Parks, Wildlife and Plant Conservation, Thailand. The colours on the map represent land use in the area. This map is modified from Sutummawong (2017)

### Reference

Manichaikul, Ani, Josyf C Mychaleckyj, Stephen S Rich, Kathy Daly, Michèle Sale, and Wei-Min Chen. 2010. 'Robust relationship inference in genome-wide association studies', *Bioinformatics*, 26: 2867-73.

Marr, Melissa M., Emily Humble, Peter W. W. Lurz, Liam A. Wilson, Elspeth Milne, Katie M. Beckmann, Jeffrey Schoenebeck, Uva-Yu-Yan Fung, Andrew C. Kitchener, Kenny Kortland, Colin Edwards, and Rob Ogden. 2025. 'Genomic Insights Into Red Squirrels in Scotland Reveal

Loss of Heterozygosity Associated With Extreme Founder Effects', *Evolutionary Applications*, 18: e70072.
Sutummawong, Nantida. 2017. 'Assessing the vulnerability of Thailand's forest birds to global change', James Cook University.
